# Centrosomal P4.1-associated protein (CPAP) is a novel regulator of ESCRT pathway function during endosome maturation

**DOI:** 10.1101/2025.08.09.669489

**Authors:** Radhika Gudi, Chenthamarakshan Vasu

**Affiliations:** Department of Pharmacology and Immunology, College of Medicine, Medical University of South Carolina, Charleston, SC 29425

**Keywords:** Centrosomal P4.1-associated protein (CPAP), endosome, endocytic vesicular transport, endosomal trafficking, endosome maturation, multi-vesicular body, endosomal sorting complex required for transport (ESCRT) pathway, Rab5-to-Rab7 conversion

## Abstract

Previously, we reported that Centrosomal P4.1-associated protein (CPAP/CENPJ/SAS-4) positively regulates multivesicular body (MVB) biogenesis and endocytic vesicular transport (EVT). Here, we show that CPAP is required for Rab5-to-Rab7 conversion, and recruitment of TSG101 to early endosome (EE) is the molecular mechanism by which CPAP promotes MVB formation. CPAP depletion disrupts Rab7 and TSG101 recruitment to EE and blocks endosome maturation. While endogenous CPAP co-precipitates with ESCRT-proteins TSG101, HRS and ALIX, exogenously expressed CPAP only interacts with TSG101. CPAP localizes to endosome and colocalizes with HRS and TSG101 during EVT progression. TSG101 recruitment to endosome, Rab5-to-Rab7 conversion, and EVT of EGFR to MVB in CPAP-depleted cells are restored by re-introduction of CPAP or overexpression of, but not endogenous, HRS. These observations show that CPAP is an ESCRT-associated protein, which functions upstream of TSG101, but in parallel to HRS, during MVB formation and EVT of cargo to the lysosome.

**Highlights:**

- CPAP-deficiency prevents Rab5-Rab7 conversion and TSG101 recruitment to early endosomes during MVB formation
- CPAP is an essential ESCRT protein
- CPAP functions parallel to HRS during MVB formation
- CPAP-TSG101 interaction is essential for Rab5-Rab7 conversion and endosome maturation

## Introduction

The endosomal trafficking/endocytic vesicular transport (EVT) pathway involves precisely coordinated signaling events that originate at the endosomes and post-endocytosis at the plasma membrane. This pathway is critical for inter-and intra-cellular communication as well as to regulate the abundance and function of proteins including cell surface receptors [1–6]. Upon endocytosis, internalized cargo such as ligand-bound epidermal growth factor receptor (EGFR) at the plasma membrane, is first routed to the early endosomes (EE). These endosomes serve as a sorting hub for endocytosed cargo and play a major role in deciding its fate. Endocytosed cargo can undergo recycling back to the surface, degradation in the lysosome, and/or release in exosomes[1, 7–9]. Formation of an intraluminal vesicle (ILV) packed intermediate endocytic organelle, referred to as the multi-vesicular body (MVB) or the late endosome (LE) [10, 11], and the sequential acquisition and activation of GTPases Rab5 and Rab7 on to EE, together coordinate endosome maturation [12–14] and precedes lysosome targeting of the endocytosed cargo.

Rab5-to-Rab7 conversion, a process involving the recruitment of Rab7 onto Rab5-positive vesicles and replacement of Rab5 with Rab7, is an important step that marks the maturation of EE to MVBs or LE [15, 16]. ILVs, essential component of MVBs, are generated in an endosomal sorting complex for recruitment and transport (ESCRT) pathway-dependent manner. ESCRT is composed of several sub-complexes: ESCRT-0, ESCRT-I, ESCRT-II and ESCRT-III. Among these ESCRT complexes, HRS (ESCRT-0) and TSG101 (ESCRT-I) are the initiators of MVB biogenesis [17–19]. In addition, several accessory proteins such as ALIX, LIP5 and ATPases such as VPS4B are also essential for the function of ESCRT pathway [20]. However, although both are essential for EVT to proceed unhindered, whether Rab5-to-Rab7 conversion and ESCRT function are sequential or parallel events and if these pathways coordinate during cargo transport is largely unknown.

Previously, we have reported the novel, positive regulatory function of CPAP (Centrosomal P4.1-associated protein; expressed by CENPJ gene), also called as SAS-4, a microtubule/tubulin-binding essential centriole biogenesis protein [21–24] during EVT of internalized EGFR cargo [25]. Prior to this report, CPAP function has been primarily attributed to its role during centriole duplication, specifically restricting the centriole length, and ciliogenesis [21, 24, 26–30]. CPAP regulates microtubule growth to produce centrioles of optimum dimension [31]. Employing gain- and loss-of-function approaches as well as the ligand-bound EGFR intracellular-trafficking model, we demonstrated that CPAP is required for efficient transport of internalized EGFR to MVB and lysosome for degradation. Depletion of CPAP resulted in disruption of MVB formation and lysosome targeting of ligand-bound EGFR, causing increased cellular abundance and constitutive signaling by this receptor [25]. Importantly, higher EGFR levels and defective EVT function due to CPAP-deficiency were correlated with increased tumorigenicity of oral cancer cells [32]. These reports demonstrated that CPAP has a positive regulatory role in MVB biogenesis and EVT of receptor cargo for degradation.

Here, we have investigated the potential molecular mechanism(s) by which CPAP positively regulates EVT. We found that both Rab5-to-Rab7 conversion and the ESCRT pathway function, which together facilitate endosome maturation, ILV and MVB biogenesis, are compromised in CPAP-depleted cells. Specifically, we have identified an interaction between CPAP and the ESCRT-I protein TSG101, and this functional interaction is more profound during endocytic trafficking. Interestingly, CPAP not only localizes to the endosome, co-localizes with ESCRT-0 protein HRS and TSG101, but it is also required for recruiting TSG101 to the endosomes during ligand-induced EVT of EGFR. Overexpression of, but not endogenous, HRS compensates for CPAP function, in terms of TSG101 recruitment to EE. Reciprocally, CPAP overexpression compensates for HRS function and restores TSG101 recruitment to EE and endosome maturation. On the other hand, TSG101 overexpression alone did not override the requirement of CPAP for endosome maturation. Our data suggests that CPAP functions in parallel to HRS in terms of TSG101 recruitment to EE, Rab5-Rab7 conversion, endosome maturation, and EVT of EGFR. Our study shows that CPAP is a critical component of the ESCRT pathway, and it functions as ESCRT-0 on the early endosome during MVB biogenesis.

## Results

### CPAP-depletion does not affect Rab5 recruitment to EE and ligand-bound EGFR positive endosomes

Our previous report [25] showed a novel function for CPAP in positively regulating EVT. Using EGFR as a model to study EVT and lysosomal targeting of endocytosed receptor, we demonstrated that MVB biogenesis and the late endosome sorting of cargo targeted for lysosomal degradation, as indicated by the lack of localization of internalized EGFR in MVBs, are defective in CPAP-deficient cells. However, recycling pathway was not impacted by CPAP deficiency. We found that CPAP depletion resulted in increased cellular levels of total and phospho-EGFR suggesting sustained EGFR signaling, and reintroduction of CPAP expression restored EVT and EGFR homeostasis. Therefore, here, we sought to understand the mechanism by which CPAP impacts the endosomal trafficking pathway, by specifically addressing how the transport of EGFR between early and late endosomes is regulated.

Rab5 and Rab7, small molecular weight guanine triphosphatases (GTPases) that function in the tethering/docking of vesicles to their target compartment, are sequentially recruited to the EE. This temporally regulated recruitment plays a critical role in endosome maturation and the transport of endocytosed receptor cargo towards the late endosomes/lysosomes for degradation [15]. Therefore, to begin to understand why CPAP-depleted cells fail to form MVB and target the receptor cargo to late endosomes, we examined if cellular levels of Rab5 and Rab7 proteins are affected upon CPAP depletion. Immunoblotting (IB) assay of HeLa cells stably expressing control-shRNA and CPAP-specific shRNA (CPAP-knockdown) showed comparable levels of Rab5 and Rab7 proteins **(Supplemental Fig. 1)** suggesting that CPAP depletion has no impact on the cellular levels of these Rab proteins. Then, using Alexa fluor 555-linked EGF (AF555-EGF), we studied if recruitment of Rab5 to endosomes and the trafficking of ligand-bound EGFR to Rab5-positive endosomes are impacted upon CPAP depletion. Confocal microscopy analysis showed, upon EGF treatment, comparable degree of Rab5 recruitment to endosomes and colocalization with EEA1- and CD63-positive endosomes in both control- and CPAP-depleted cells **(Fig. 1A&B)**. Furthermore, both maximum projection and single Z-scan images showed that colocalization of internalized AF555-EGF with Rab5 was not considerably different at the two time-points tested **(Fig. 1C)**, suggesting that CPAP depletion does not affect Rab5 recruitment or early stages of EVT pathway. Of note, confocal microscopy images showed no difference in the relative fluorescence intensity of Rab5 between the control and CPAP-depleted cells confirming that Rab5 levels or function are not influenced by CPAP.

**Fig. 1.**
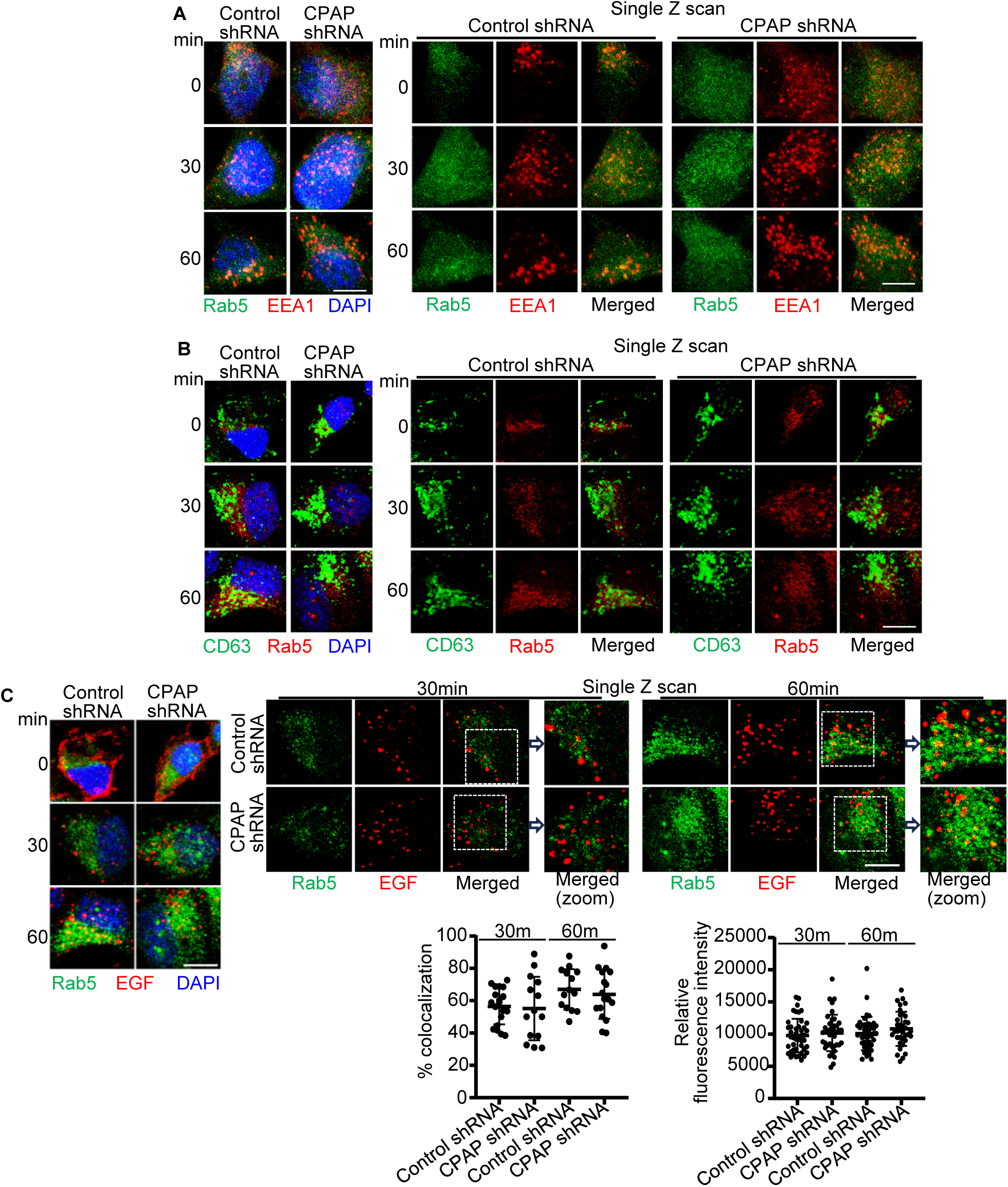
CPAP-depletion does not affect Rab5 recruitment to endosomes and ligand-bound EGFR positive endosomes. HeLa cells expressing control or CPAP shRNA were treated with AF555-EGF ligand for indicated time points and stained for Rab5 and EEA1 (A), Rab5 and CD63 (B), or Rab5 (along with AF555-EGF) and imaged by confocal microscopy (C). Left panels (A-C): maximum intensity-projection images of confocal Z stacks. Right panels (A-C): Single Z-plane of images. Bottom left graph in C: colocalization (yellow) was quantified by counting percentages of EGF-positive (red) puncta containing Rab5 (green) puncta in representative single Z-planes of each cell and quantified from multiple cells across at least 3 experiments. Bottom right graph in C; Relative integrated fluorescence intensity values of Rab5 staining quantified in representative single Z-planes of each cell and quantified from multiple cells across at least 3 experiments. Scale bar: 10µm. *p*-values were not statistically significant. Zoomed images correspond to the dashed inset boxes of the indicated images.

### CPAP-depletion disrupts Rab7 recruitment to endosomes and ligand-bound EGFR positive endosomes

During endosome maturation and cargo sorting for MVB formation, Rab7 is recruited to the maturing endosome and replaces Rab5 (referred to as the Rab5-to-Rab7 conversion) in a highly coordinated manner [15]. Therefore, we determined if Rab7 recruitment to endosome and ligand-bound EGFR localization on Rab7-positive endosomes are dysregulated in CPAP-depleted cells. Control and CPAP-depleted HeLa cells were examined for the localization of Rab7 on early and late endosomes upon EGF treatment. We found that Rab7 colocalization with EEA1 and CD63 positive endosomes was significantly lower in CPAP-depleted cells compared to control cells **(Fig. 2A&B)**. Cells treated with AF555-EGF were also stained with Rab7 to determine if the ligand bound EGFR cargo is transported to Rab7-positive late endosomes. Both maximum projection and single Z-scan images show that ligand bound EGFR localization on Rab7-positive endosomes was significantly lower in CPAP-depleted cells compared to control cells, at both time points tested **(Fig. 2C)**. Furthermore, perinuclear clustering of Rab7 was visibly diminished in these cells upon CPAP depletion. Importantly, although the IB assay showed cellular expression levels of Rab7 were unaffected by CPAP depletion **(Supplemental Fig. 1),** the relative fluorescence intensity of Rab7 in CPAP-depleted cells after ligand binding was significantly lower, suggesting its reduced clustering on endosomes during EVT progression **(Fig. 2C).** These observations suggest a defect in the activation and recruitment of Rab7 to the endosomes under CPAP deficiency. These results also suggest that our observation of failed transport of ligand bound EGFR to the MVBs and lysosomes [25] is, at least in part, due to the failure in recruiting Rab7 to endosomes, resulting in diminished endosome maturation.

**Fig. 2.**
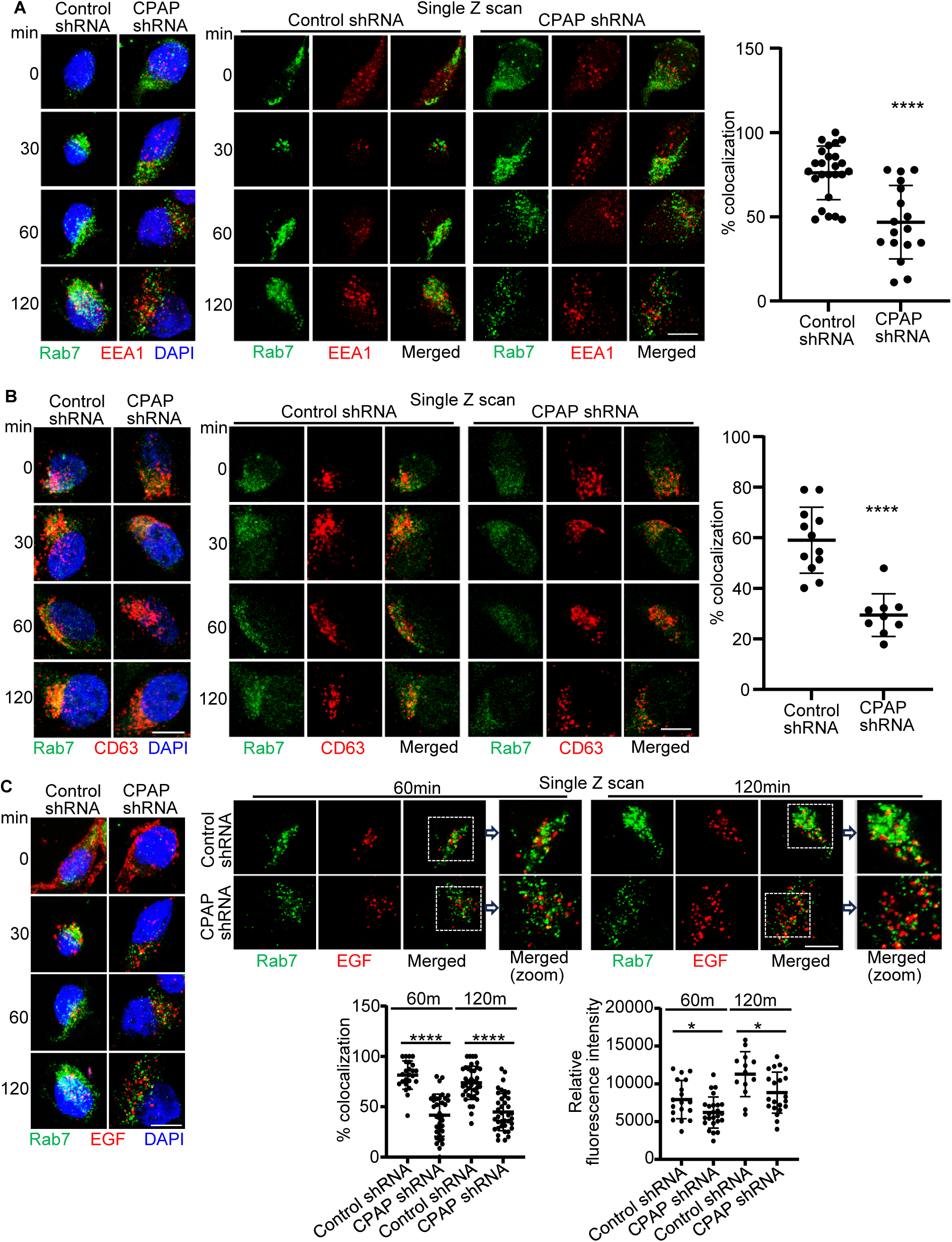
CPAP-depletion disrupts Rab7 recruitment to endosomes and ligand-bound EGFR positive endosomes. HeLa cells expressing control or CPAP shRNA were treated with AF555-EGF ligand for indicated time points and stained for Rab7 and EEA1 (A), Rab7 and CD63 (B), or Rab7 (along with AF555-EGF) and imaged by confocal microscopy (C). Left panels (A-C): maximum intensity-projection images of confocal Z stacks. Middle panels of A&B and right panel of C: Single Z-plane of images. Right panel in A: colocalization (yellow) was quantified by counting percentages of EEA1-positive (red) puncta containing Rab7 (green) puncta in representative single Z-planes of each cell and quantified from multiple cells from 60 min timepoint of one representative experiment. Right panel in B: colocalization (yellow) was quantified by counting percentages of EGF-positive (red) puncta containing Rab7 (green) puncta in representative single Z-planes of each cell and quantified from multiple cells from 60 min timepoint of one representative experiment. Object-based colocalization macro tool of FIJI was employed. Bottom left graph in C: colocalization (yellow) was quantified by counting percentages of EGF-positive (red) puncta containing Rab7 (green) puncta in representative single Z-planes of each cell and quantified from multiple cells across at least 3 experiments. Bottom right graph in C; Relative integrated fluorescence intensity values of Rab7 staining quantified in representative single Z-planes of each cell and quantified from multiple cells across at least 3 experiments. Zoomed images correspond to the dashed inset boxes of the indicated images. Scale bar: 10µm. *p*-values: *<0.05, ****<0.0001 by unpaired non-parametric Mann-Whitney test.

### CPAP depletion disrupts Rab5-to-Rab7 conversion

Since we found that Rab7 recruitment to endosome is defective in CPAP-depleted cells, we used Airyscan super-resolution microscopy to determine if receptor internalization-induced endosomal colocalization of Rab5 and Rab7 is impacted when CPAP is depleted. We employed both shRNA and siRNA approaches **(Supplemental Figs. 1&2)** that were validated in our previous reports [25, 32]. Control and CPAP-depleted HeLa cells were treated with untagged EGF, stained, and imaged to detect Rab5 and Rab7 colocalization. Both maximum intensity projection and single Z images showed that colocalization of Rab7 with Rab5 was significantly lower in CPAP-depleted cells **(Fig. 3A and Supplemental Fig. 3)**, indicating that Rab7 recruitment to Rab5-positive endosomes and Rab5-to-Rab7 conversion is disrupted in the absence of CPAP. To validate this observation, control and CPAP-depleted HeLa cells were transfected with GFP-Rab5 and mCherry-Rab7 vector constructs, treated with EGF to induce EVT, and imaged to detect GFP-mCherry (Rab5-Rab7) colocalization. **Fig. 3B** shows that Rab7 localization to Rab5-positive endosome is significantly lower in CPAP-depleted cells compared to control cells. Since CPAP-depleted cells show diminished recruitment of Rab7 to endosomes during EGFR trafficking and lower overall fluorescence-staining intensity for Rab7 **(Fig. 2)**, we examined if overexpression of Rab7 restores endosome maturation and the EVT of endocytosed EGFR to MVBs. Control and CPAP-depleted HeLa cells were transfected with vector constructs encoding cDNA for GFP-Rab7, either wild-type (WT) or dominant negative (DN) and subjected to EGFR trafficking assay using AF555-EGF, followed by confocal imaging. CPAP-depleted cells, as expected, showed a lower degree of localization of ligand-bound EGFR to the CD63-positive late endosomes/MVBs compared to control cells **(Fig. 3C)**. Importantly, overexpression of WT Rab7 restored EGFR trafficking to MVB in CPAP-depleted cells as indicated by the localization of AF555-EGF to the CD63-positive endosomes. However, exogenous expression of DN Rab7 not only failed to restore the EVT of EGFR, but also suppressed the MVB localization of AF555-EGF in control cells. These results substantiate an important role for Rab7 recruitment to endosome in CPAP-mediated positive regulation of EVT and trafficking of ligand-bound EGFR to MVB. Further, this data also suggests that CPAP functions at the level of, by direct interaction, or upstream of Rab7 recruitment to facilitate EVT. In this regard, we conducted an immunoprecipitation (IP) assay using lysates of GFP-, GFP-Rab5- or GFP-Rab7-expressing cells to determine if Rab5 and Rab7 interact with CPAP. Immunoblotting of GFP-specific pulldowns suggested no interactions between CPAP and these Rab proteins **(Supplemental Fig. 4)**. This lack of CPAP-Rab7 interaction suggests that although endosome maturation and Rab7 recruitment to the endosome is affected upon CPAP depletion, another mechanism or player that operates in parallel or upstream of Rab5-to-Rab7 conversion could be the target of CPAP-mediated positive regulation of EVT.

**Fig. 3.**
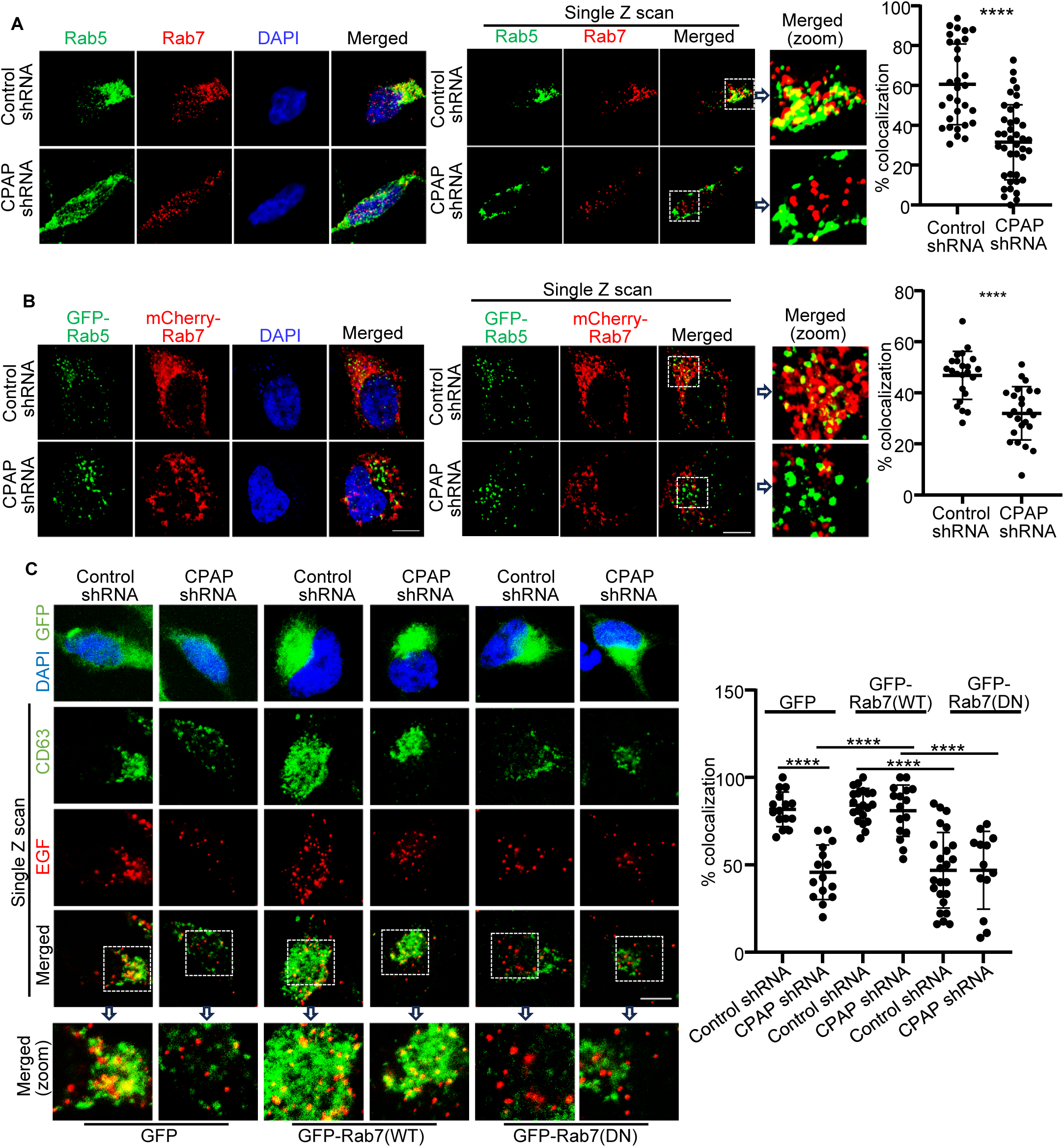
CPAP depletion disrupts Rab5-to-Rab7 conversion. A) HeLa cells expressing control or CPAP shRNA were treated with untagged EGF for 60 min, stained for Rab5 and Rab7, and imaged by Airyscan super-resolution microscopy. Left panel: maximum intensity-projection images of Z stacks; middle panel: single Z-plane of images; right panel: colocalization (yellow) was quantified by counting percentages of Rab5-positive (green) puncta containing Rab7 (red) puncta in representative single Z-planes of each cell and quantified from multiple cells across at least 3 experiments. B) HeLa cells expressing control or CPAP shRNA were transfected with GFP-Rab5 and mCherry-Rab7 constructs, treated with untagged EGF for 60 min, and imaged by Airyscan super-resolution microscopy. Left panel: maximum intensity-projection images of Z stacks; middle panel: single Z-plane of images; right panel: colocalization (yellow) was quantified by counting percentages of GFP-Rab5-positive (green) puncta containing mCherry-Rab7 (red) puncta in representative single Z-planes of each cell and quantified from multiple cells across at least 3 experiments. Object-based colocalization macro tool of FIJI was employed. C) HeLa cells stably expressing control or CPAP-shRNA were transfected with control vector (GFP), or GFP-Rab7 (WT; wild-type) or GFP-Rab7 (DN; dominant negative) vector constructs, treated with AF555-EGF ligand for 60 min, and stained for CD63 to mark MVBs/late endosomes, and imaged by confocal microscopy to determine AF555-EGF and CD63 colocalization. Left top row: GFP expression in control and Rab7 construct-expressing cells; left lower rows: representative single Z-plane of images showing ligand-bound EGFR-CD63 colocalization; right panel: colocalization (yellow) was quantified by counting percentages of EGF-positive (red) puncta containing CD63 (green) puncta in representative single Z-planes of each cell and quantified from multiple cells across at least 3 experiments. Zoomed images correspond to the dashed inset boxes of the indicated images. Scale bar: 10µm. *p*-values: ****<0.0001 by unpaired non-parametric Mann-Whitney test.

### Recruitment of TSG101, an ESCRT-I component, to the endosomes is compromised in CPAP-depleted cells

Endosome maturation is concomitantly regulated by the Rab GTPases and ESCRT complexes, especially during the early to late endosome transition. ESCRT proteins (0, I, II, III and its accessory counterparts) are sequentially recruited to the EE membrane and they drive the inward bending of cargo-associated membrane to generate intraluminal vesicles (ILVs) that form the mature MVB [20, 33]. These ILV-rich MVBs fuse with lysosomes to form endolysosomes and degrade the endocytosed receptor cargo. Importantly, ESCRT disruption has been reported to inhibit the completion of Rab5-to-Rab7 conversion [34]. Further, endocytic cargo corralling by ESCRTs upstream of Rab5-Rab7 conversion has been hypothesized in a recent study [35]. Nevertheless, coordination of ESCRT and Rab5-to-Rab7 conversion pathways during endosome maturation and MVB formation is not understood. Here, we tested the dynamics of ESCRT function during the EVT of EGFR in control- and CPAP-depleted HeLa cells. To determine ESCRT recruitment to endosomes, EGF-treated cells were stained using HRS (ESCRT-0), TSG101 (ESCRT-I) and ALIX (ESCRT accessory component) -specific antibodies together with the anti-EEA1 antibody and subjected to Airyscan super-resolution microscopy. Of note, specificities of the ESCRT component antibodies were confirmed by immunoblotting and immunofluorescence of control- and -specific siRNA treated cells **(Supplemental Fig. 5A&B)**. Importantly, we found that the recruitment of HRS or ALIX to the EEA1-positive endosomes, as assessed by quantifying the number of colocalized puncta, was not different between control-and CPAP-depleted cells **(Fig. 4)**. However, the localization of TSG101, an ESCRT-I component, which triggers MVB formation and downstream EVT events [18], on the EE was significantly lower in CPAP-depleted cells when compared to control cells, suggesting that CPAP may be required for TSG101 recruitment to the endosome. Importantly, it has been shown before that ESCRT-0 protein HRS triggers TSG101 localization/recruitment to the endosomes [17]. Our experiment using HeLa cells to re-assess this aspect showed that TSG101 recruitment to the EE during EGF treatment is, in fact, diminished severely in HRS-depleted cells **(Supplemental Fig. 6)**. These observations suggest that CPAP and HRS function similarly, in terms of TSG101 recruitment during EVT.

**Fig. 4.**
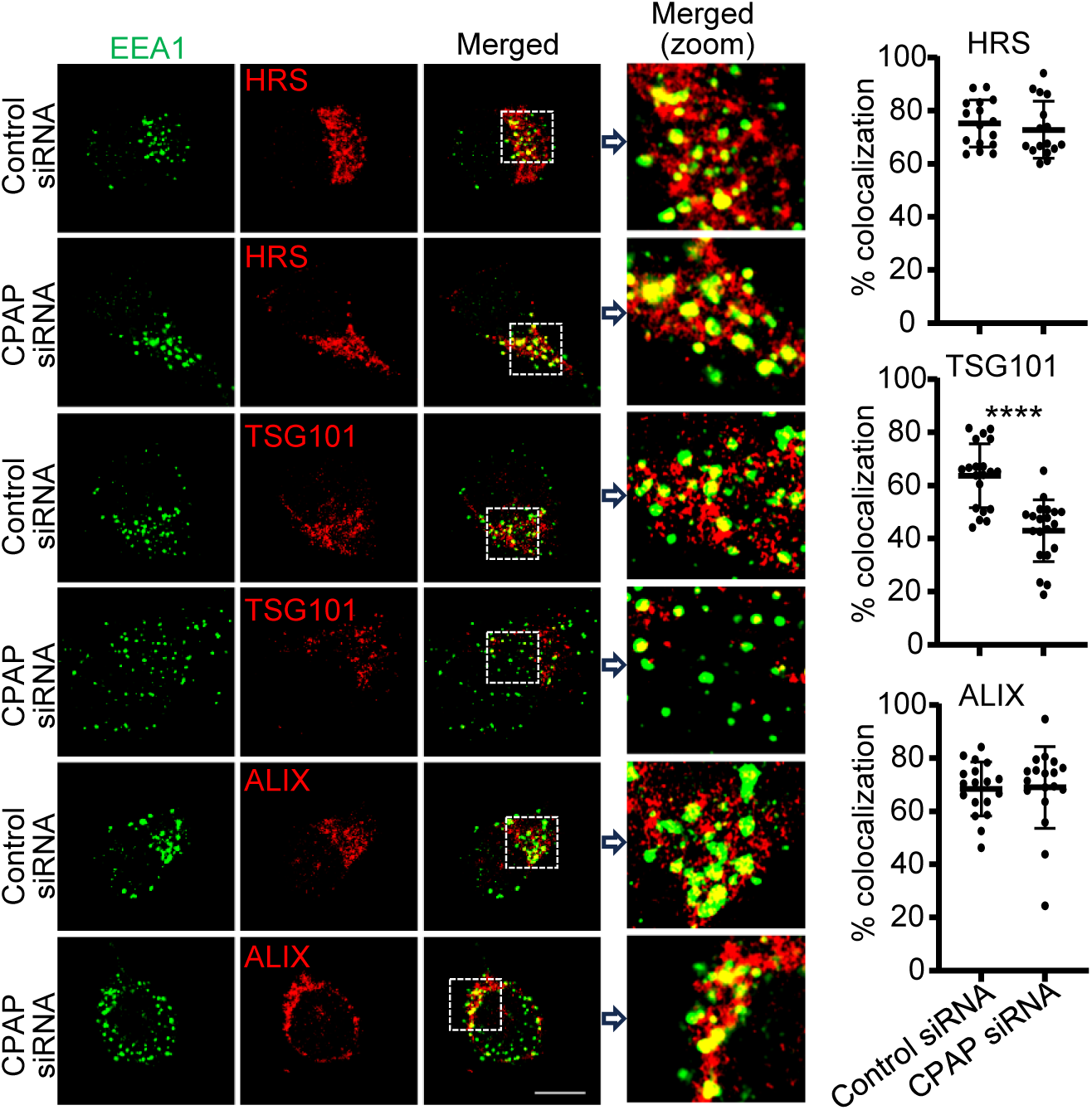
Recruitment of TSG101, an ESCRT-I component, to the endosomes is compromised in CPAP-depleted cells. HeLa cells expressing control or CPAP shRNA were treated with untagged EGF for 60 min, stained for ESCRT pathway proteins HRS, TSG101 or ALIX along with EEA1 (to mark EE), and imaged by Airyscan super-resolution microscopy. Left panels: representative single Z-plane of images showing localization of HRS, TSG101, or ALIX on EEA1- positive puncta. Zoomed images correspond to the dashed inset boxes of the indicated images. Right panels: colocalization (yellow) was quantified by counting percentages of EEA1-positive (green) puncta containing ESCRT protein-positive (red) puncta in representative single Z-planes of each cell and quantified from multiple cells across at least 3 experiments. Scale bar: 10µm. *p*-values: ****<0.0001 by unpaired non-parametric Mann-Whitney test.

### CPAP is detected on endosomes in quiescent and EGF-treated cells and it colocalization with ESCRT proteins increases during EVT

Since CPAP depletion appears to disrupt TSG101 recruitment to the EE, employing different approaches and multiple EEA1 antibodies, we examined if CPAP localizes to EE under quiescent state and upon EVT progression. Control- and CPAP-depleted cells were left untreated or treated with EGF and subjected to immunostaining for CPAP and EEA1 followed by confocal microscopy. As observed in **Supplemental Fig. 7**, considerable proportion of EEA1-positive endosomes were also positive for CPAP staining in quiescent and EGF stimulated cells. Fluorescence intensity of CPAP on EEA1-positive endosomes was significantly diminished in the CPAP-depleted cells, emphasizing the specificity of endosomal staining of CPAP. As observed in **Fig. 5A**, CPAP colocalizes with EEA1-positive endosomes in untreated cells, and shows a significant time-dependent increase in this colocalization during EVT progression in EGF-treated cells. We then examined if CPAP colocalizes with TSG101 and HRS on EE. Control and EGF treated cells were stained for EEA1 and CPAP, alone or along with HRS and TSG101, and subjected to super-resolution microscopy. Examination of tri-protein colocalization between CPAP and HRS or CPAP and TSG101, on EEA1-positive endosomes, also revealed that CPAP along with these ESCRT proteins localizes to the EE during EVT progression. While CPAP and ESCRT proteins (HRS and TSG101) were found to be colocalized to a small number of EEA1- positive endosomes in untreated cells, this tri-protein colocalization showed a time-dependent increase in EGF-treated cells **(Fig. 5B)**. Overall, these observations suggest that CPAP is recruited to the EE, along with ESCRT-0 and ESCRT-I proteins, during EVT of cell surface receptor cargo.

**Fig. 5.**
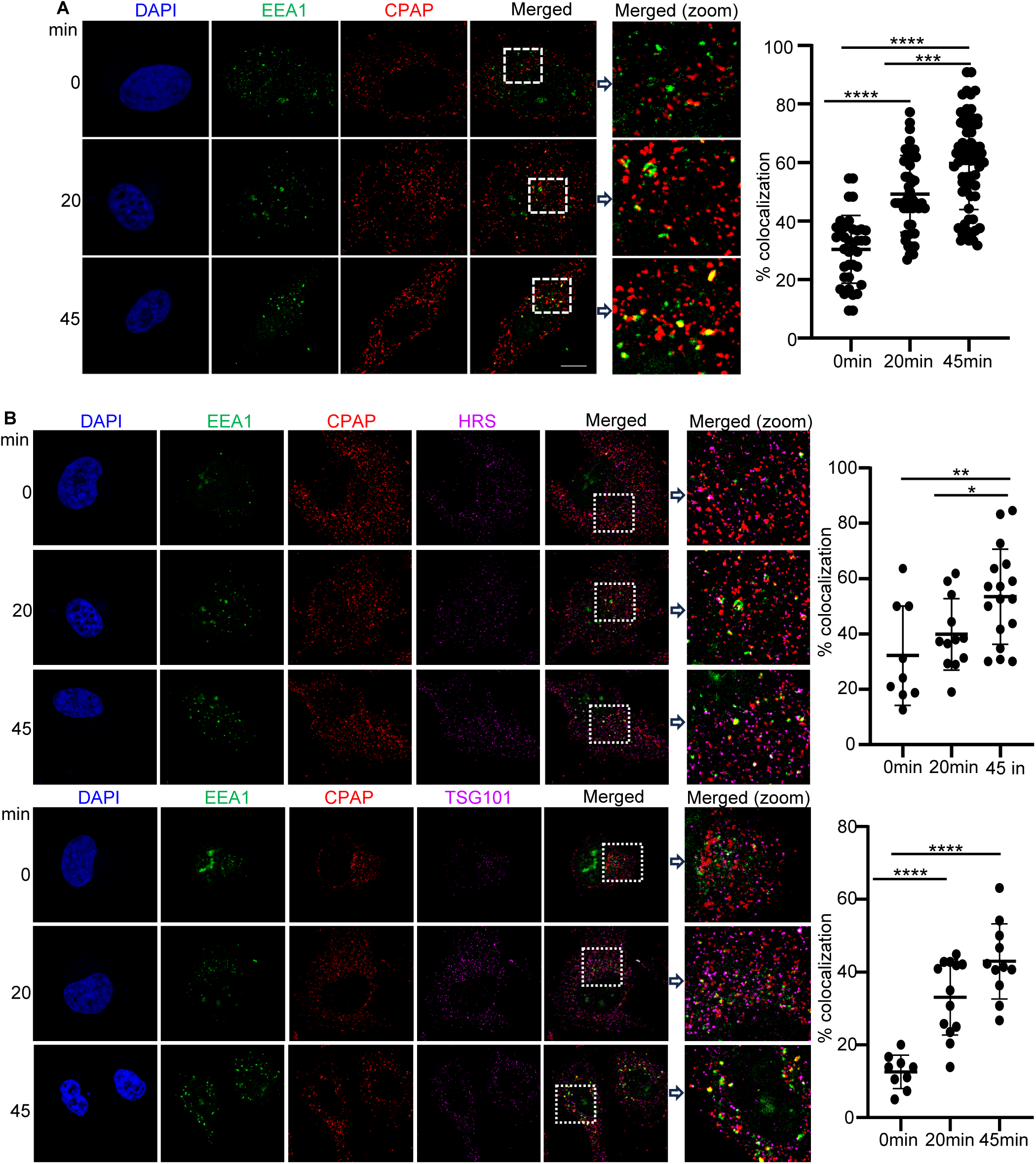
CPAP is detected on endosomes in quiescent and EGF-treated cells and its colocalization with ESCRT proteins increases during EVT. A) HeLa cells were treated with untagged EGF for different time-points, stained for EEA1 and CPAP, and imaged by Airyscan super-resolution microscopy. Left panel: single Z-plane of images showing EEA1 and CPAP staining of representative cells; right panel: colocalization (yellow) was quantified by counting percentages of EEA1-positive (green) puncta containing CPAP (red) puncta in representative single Z-planes of each cell and quantified from multiple cells across at least 3 experiments. B) HeLa cells treated with untagged EGF for indicated time points were stained with AF488-linked sheep anti-EEA antibody, rabbit anti-CPAP, and mouse anti-HRS or -TSG101 primary antibodies followed by AF647- and AF568-linked anti-rabbit and -mouse secondary antibodies and imaged using Lightning super-resolution microscopy. Left panel: representative single Z-plane of images showing colocalization of CPAP along with HRS or TSG101 on EEA1 positive puncta. Right panels: triple colocalization (white) was quantified by counting percentages of EEA1-positive (green) puncta containing CPAP (red) and HRS/TSG101 (magenta) puncta in representative single Z-planes of each cell and quantified from multiple cells across at least 3 experiments. Zoomed images correspond to the dashed inset boxes of the indicated images. Scale bar: 10µm. *p*-values: *<0.05, **<0.01, ***<0.001, ****<0.0001 by unpaired non-parametric Mann-Whitney test.

To substantiate the observation that CPAP localizes to endosomes and is actively recruited to endosomes during EVT progression, we have examined the presence of CPAP on EEA1-positive structures by transmission electron microscopy (TEM). TEM images of EGF-treated cell sections that were labeled using CPAP-specific antibody, followed by gold particle-linked secondary antibody showed positive-labeling of endosomes for CPAP **(Supplemental Fig. 8A&B**). IB of endosome/vesicle-rich fraction from CPAP-overexpressing (GFP-CPAP) cells showed the presence of CPAP along with Rab5 and Rab7 **(Supplemental Fig. 9A)**. Further, the vesicle-rich fraction of non-transfected, EGF-treated cells showed higher levels of endogenous CPAP compared to that of untreated cells **(Supplemental Fig. 9B)**. Collectively, these observations indicate the presence of CPAP on endosomes and its vesicular levels are higher during EVT progression.

### Exogenous expression of CPAP restores TSG101 recruitment on to EE in CPAP and HRS-depleted cells

Since we observed that CPAP can localize to the EE, it actively colocalizes with ESCRT proteins on EE during EVT progression, and its depletion disrupts TSG101 recruitment to the EE, we next determined if reintroduction of CPAP expression can restore TSG101 recruitment to the endosome in CPAP- and HRS-knockdown cells. HeLa cells stably transfected with doxycycline-inducible, RNAi-resistant, GFP-tagged-CPAP cDNA were treated with CPAP-specific-siRNA to deplete endogenous CPAP expression, induced exogenous GFP-CPAP expression, treated with EGF to induce EVT and subjected to immunostaining with the anti-EEA1- and -TSG101 antibodies for Airyscan-based super-resolution microscopy as depicted in **Fig. 6A**. As expected, CPAP-siRNA and HRS-siRNA treated control cells showed significantly lower colocalization of TSG101 and EEA1 compared to control siRNA treated cells. However, upon induction of GFP-CPAP expression in CPAP-siRNA treated cells, localization of TSG101 on EEA1 positive endosomes was significantly increased **(Fig. 6B)**, suggesting that TSG101 recruitment to endosomes is restored, at least in part, by reintroduction of CPAP. Importantly, overexpression of CPAP in HRS-depleted cells, similar to CPAP-depleted cells, restored TSG101 recruitment to the EE. Furthermore, exogenous CPAP expression caused a significant increase in the localization of ligand-bound EGFR on CD63-postive endosomes (MVBs) in CPAP- and HRS-depleted cells suggesting restoration of endosome maturation **(Supplemental Fig. 10)**, reiterating that CPAP functions similarly to, and compensates for, HRS during EVT.

**Fig. 6.**
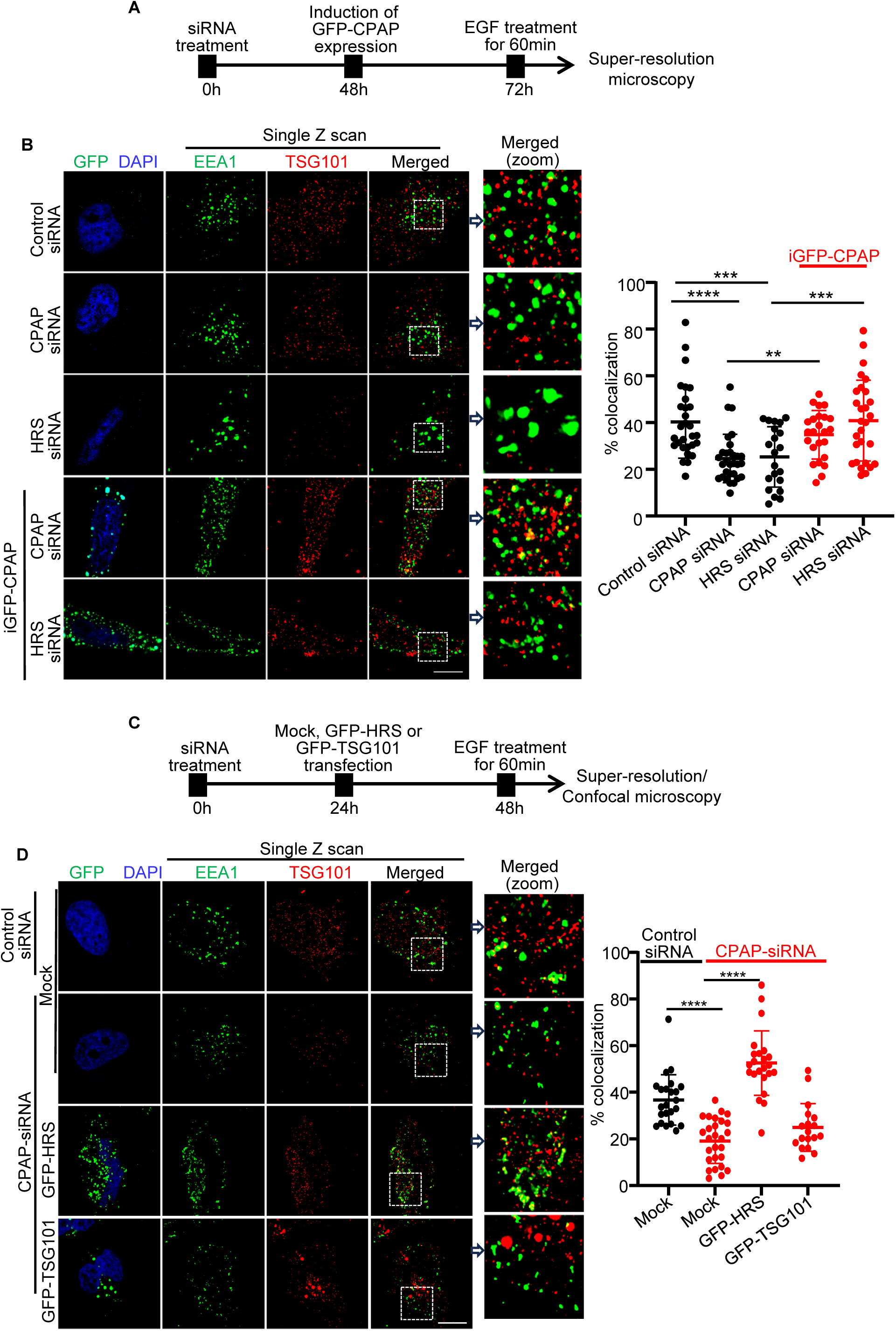
Exogenous expression of HRS and CPAP restore TSG101 recruitment on to EE in CPAP and HRS-depleted cells respectively. A) Schematic depiction of experimental strategy using control-, CPAP- and HRS-specific siRNA treated HeLa cells with and without siRNA-resistant GFP-CPAP expression. B) Lightning super resolution microscopy images showing cells stained for EEA1 and TSG101. C) Schematic depiction of experimental strategy using control-and CPAP-specific-siRNA treated HeLa cells with and without GFP-HRS or GFP-TSG101 expression. D) Lightning super resolution microscopy images showing cells stained for EEA1 and TSG101. Left panels of B and D: representative single Z-plane of images showing localization of TSG101 on EEA1 positive puncta in cells with and without GFP expression. Right panels of B and D: colocalization (yellow) was quantified by counting percentages of EEA1-positive (green) puncta containing TSG101-positive (red) puncta in representative single Z-planes of each cell and quantified from multiple cells across at least 3 experiments. Zoomed images correspond to the dashed inset boxes of the indicated images. Scale bar: 10µm. *p*-values: **<0.01, ***<0.001, ****<0.0001 by unpaired non-parametric Mann-Whitney test.

### HRS, but not TSG101, overexpression restored TSG101 recruitment onto endosome and EVT function in CPAP-depleted cells

It has been shown that EVT and MVB formation are facilitated by HRS-TSG101 interaction [17, 19, 36]. Since CPAP-depleted cells are defective in MVB formation and EVT of EGFR [25], we tested if overexpression of HRS or TSG101 alone was sufficient to restore TSG101 recruitment to endosomes and trafficking of EGFR to MVBs in these cells. Control- and CPAP -siRNA treated HeLa cells were subjected to GFP-TSG101 or HRS expression vector transfection **(Supplemental Fig. 11)** and examined for colocalization of TSG101 on EEA1-positive endosomes **(Fig.6C)**. As observed in **Fig. 6D**, no significant increase in the localization of TSG101 on EEA1-positive endosomes of CPAP-depleted cells, compared to mock-transfected cells, was observed upon TSG101 overexpression itself. Unlike TSG101-overexpression, HRS-overexpression in CPAP-depleted cells led to significantly higher colocalization of TSG101 on EEA1-positive endosomes as compared to mock transfected cells. This suggests that, similar to CPAP’s ability to compensate for HRS function **(Fig. 6B)**, higher protein levels of HRS can compensate for CPAP function in terms of TSG101 recruitment to the EE. Our data indicates that CPAP functions upstream of TSG101, but appears to be in parallel to HRS, in recruiting ESCRT pathway proteins to the endosome.

To validate this notion, EGFR trafficking to MVBs in CPAP-depleted cells that are overexpressing TSG101 or HRS was examined after treating with AF555-EGF. **Supplemental. Fig. 12** shows that the degree of AF555-EGF localization to CD63 positive endosomes was not significantly different in CPAP-depleted, TSG101-overexpressing cells as compared to CPAP-depleted cells. On the other hand, AF555-EGF colocalization with CD63-positive endosomes was higher in CPAP-depleted, HRS-overexpressing cells compared to CPAP-depleted controls. Overall, these results, in the context of our previous report [25], show that the ESCRT pathway function is disrupted upon CPAP depletion due to failed TSG101 recruitment to EE, leading to defective MVB formation and receptor cargo transport. These results reiterate the notion that CPAP acts upstream of TSG101 and parallel to HRS in the MVB biogenesis pathway.

### CPAP depletion does not impact the cellular levels of TSG101

Since CPAP depletion appears to disrupt TSG101 recruitment onto the endosome, we examined whether cellular levels of TSG101 and other key ESCRT proteins are altered upon CPAP depletion. First, lysates of control- or CPAP-depleted HeLa **(Fig. 7A)** and HEK293T **(Fig. 7C)** cells were subjected to immunoblotting to detect HRS, TSG101 and ALIX. Fig. 7A and 7C shows that the levels of these ESCRT proteins are comparable in control- and CPAP-depleted cells in both cell lines. Cellular levels of these ESCRT proteins appear to be unaltered in CPAP-depleted cells under EGF stimulation as well **(Supplemental Fig. 13)**. We then stained control-and CPAP-depleted HeLa and HEK293T cells to detect intracellular levels of CPAP and TSG101 and analyzed by flow cytometry. This assay also showed that cellular levels of TSG101 remain unaltered by CPAP depletion **(Fig. 7B and 7D)**. These observations, in the context of afore-described disrupted recruitment of TSG101 on to endosome in CPAP-depleted cells, suggest that CPAP positively regulates direct or HRS-dependent recruitment of TSG101 onto the endosome, but does not have an influence on the cellular levels or stability of this ESCRT protein during EVT.

**Fig. 7.**
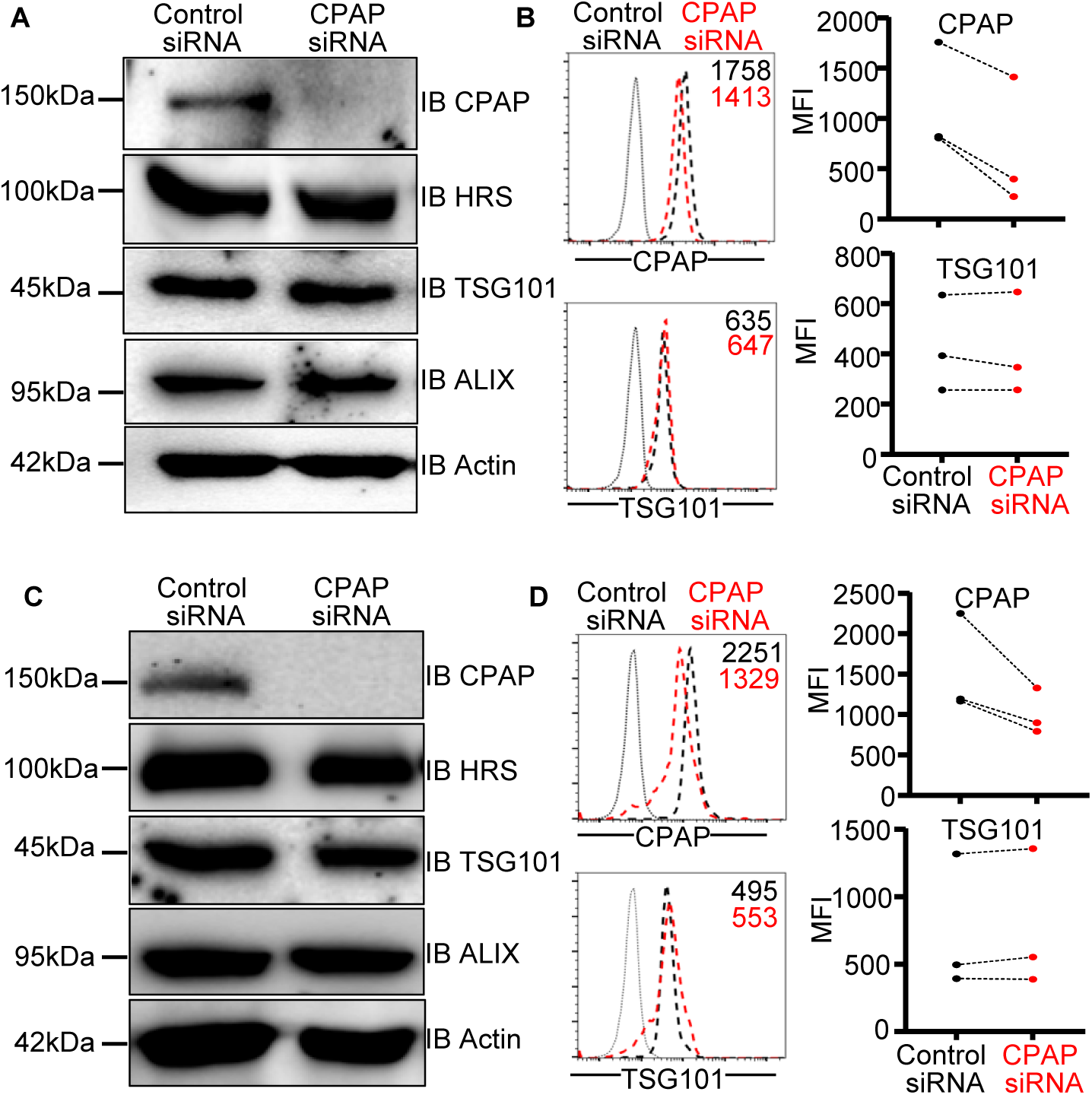
CPAP depletion does not impact the cellular levels of TSG101. HeLa and HEK293T cells were treated with control or CPAP specific siRNA for 48h and subjected to IB to detect the cellular levels of CPAP and ESCRT proteins (A&C). Cells were also subjected to intracellular staining and flow cytometry to detect cellular levels of CPAP and TSG101 (B&D). Representative IB images of three independent experiments that produced similar trends in outcomes are shown for panels A&C. Representative histogram overlay plots (left) and mean fluorescence intensity (MFI) values of three independent experiments (right) are shown for panels B&D.

### CPAP interacts with TSG101, and this interaction is enhanced upon EVT

To begin to understand how CPAP regulates TSG101 recruitment to the endosomes, we tested if the ESCRT proteins interact with CPAP. HEK293T cell lysates were subjected to IP using HRS, ALIX and TSG101 -specific antibodies separately, followed by IB to detect these proteins along with CPAP. Interestingly, CPAP was detected in the pull-down of all three ESCRT components **(Fig. 8A)** suggesting that it is associated with ESCRT machinery proteins under quiescent state. To substantiate the observed interaction between CPAP and ESCRT proteins, HEK293T cells stably transfected with inducible GFP-CPAP vector were treated with doxycycline to induce GFP-CPAP expression, left untreated or treated with EGF, and subjected to IP and IB. GFP-specific IP of untreated cells probed for ESCRT proteins showed only TSG101, but not HRS or ALIX **(Fig. 8B)**. Surprisingly, the amount of TSG101 pulled down along with GFP-CPAP was profoundly higher in EGF-stimulated cells compared to untreated cells, suggesting that the interaction between CPAP and TSG101 is amplified during the progression of EVT.

**Fig. 8.**
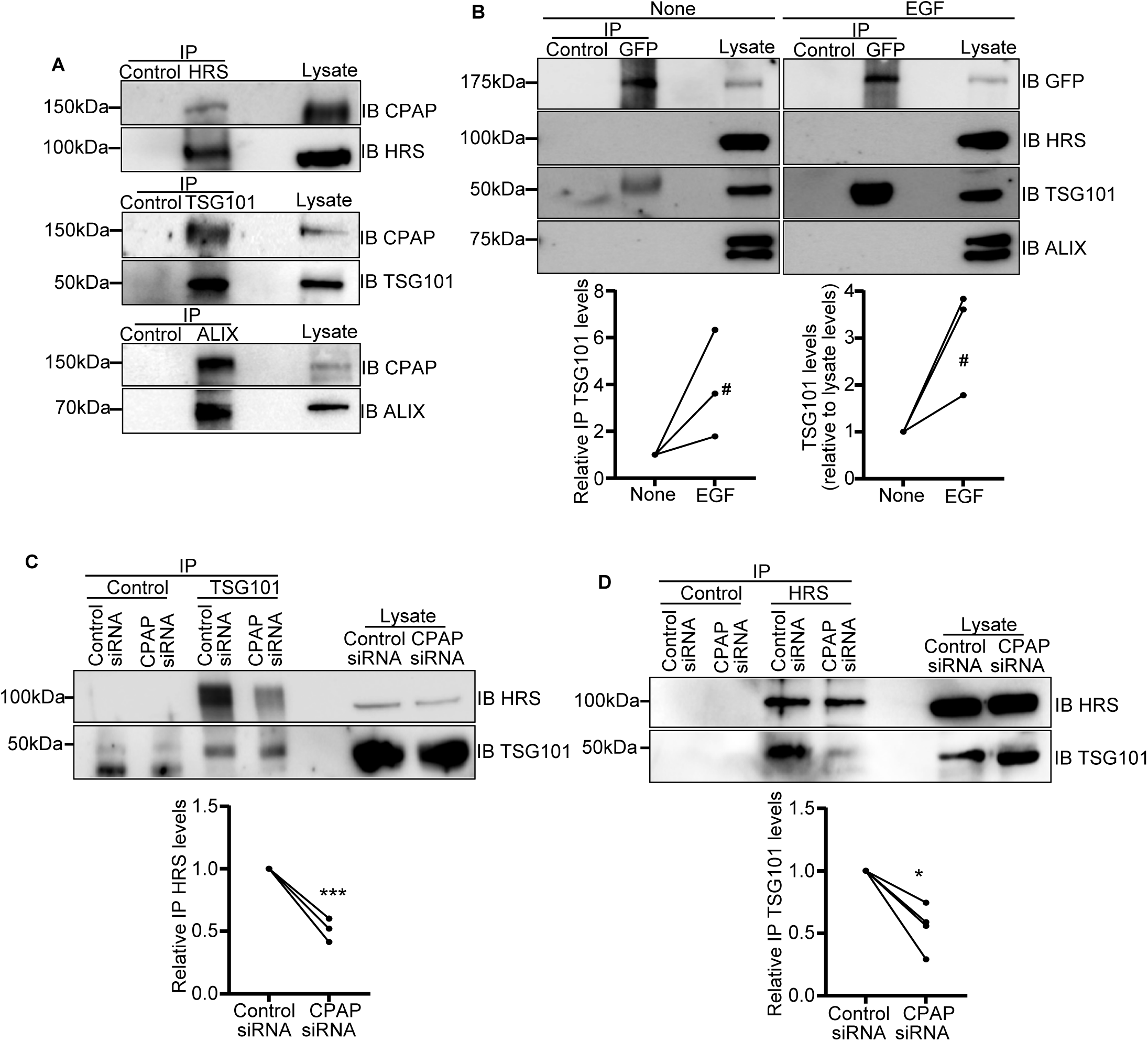
CPAP interacts with TSG101, and this interaction is enhanced upon EVT. A) Lysates of HeLa cells were subjected to IP using control antibody or anti -HRS, -TSG101- or - ALIX antibodies followed by IB using indicated antibodies. Representative of three independent experiments. B) Doxycycline-inducible GFP-CPAP expressing HEK293T cells were treated with doxycycline for 24h, left untreated or treated with EGF ligand for 45 mins, and subjected to IP using anti-GFP antibody and IB using indicated ESCRT protein-specific antibodies. Upper panel: representative IB images; lower left: relative densitometric values of TSG101 bands from three independent experiments with none as base value of 1 for each experiment; lower right: densitometric values of TSG101 bands of IP (relative to lysate bands) with none as base value of 1 for each experiment. (C&D) HeLa cells were treated with control- and CPAP-specific siRNA for 48h, and EGF for 60 min and subjected to IP using control antibody or anti-TSG101 (C) or - HRS (D) antibodies, followed by IB using TSG101- and HRS-specific antibodies. Upper panels: representative IB images; lower panels: relative densitometric values of HRS (left panel) and TSG101 (right panel) bands from three independent experiments with control siRNA as base value of 1 for each experiment. *p*-values: #<0.1, *<0.05, ***<0.001 by paired *t*-test.

### HRS-TSG101 interaction is abrogated in the absence of CPAP during EVT

To evaluate the role of CPAP in bridging ESCRT recruitment during EVT, HRS-TSG101 interaction was evaluated in EGF-treated, control or CPAP-depleted cells. While the amount of TSG101 immunoprecipitated from the control- and CPAP-depleted cells was comparable, CPAP-depleted cells revealed a significant reduction in the amount of HRS that was pulled down, as compared to control cells, by TSG101-specific antibody **(Fig. 8C)**. Reciprocal IP of EGF-treated, control- or CPAP-depleted cells using HRS antibody showed lower TSG101 protein pulldown from CPAP-depleted cells compared to control cells **(Fig. 8D)**. These observations suggest that CPAP is essential for the functional interaction between HRS and TSG101 and the recruitment of TSG101 to endosome by HRS during endocytosed receptor cargo transport.

### Rab5-to-Rab7 conversion and EGFR trafficking to endosome are restored in CPAP-depleted cells upon HRS, but not TSG101, overexpression

Our previous study showed that lysosomal targeting of ligand-bound EGFR to lysosome is disrupted upon CPAP depletion [25]. Our results here show that, although CPAP does not interact with or affect the cellular levels of Rab5 or Rab7, Rab5-to-Rab7 conversion is blocked in CPAP-depleted cells **(Fig. 3 and Supplemental Figs 1&3)**. Further, our observations that CPAP not only interacts with TSG101, but also recruits TSG101 to endosomes and restores EVT function in HRS-depleted cells **(Figs. 6&8)** suggest that disruption of ESCRT pathway function is responsible for the defective Rab5-to-Rab7 conversion in CPAP-depleted cells. Therefore, first we tested if overexpression of ESCRT components, TSG101 or HRS, is sufficient to restore Rab5-to-Rab7 conversion. Control- and CPAP-siRNA-expressing HeLa cells were subjected to GFP-TSG101 or GFP-HRS expression vector transfection and examined for Rab5 and Rab7 colocalization in them using Lightning super-resolution microscopy as shown in **Fig. 9A**. As observed in **Fig. 9B**, overexpression of TSG101 in CPAP-depleted cells did not increase colocalization of Rab5 and Rab7 as compared to control cells. On the other hand, HRS overexpression in CPAP-depleted cells increased the localization of Rab7 onto the Rab5-positive endosomes as compared to control cells. Next, we examined whether the EVT function and localization of internalized ligand-bound EGFR to Rab7-positive endosomes are restored in CPAP-depleted cells upon overexpression of TSG101 or HRS. As shown in **Fig. 9C**, HRS-overexpression in CPAP-depleted cells caused an increase in the localization of ligand (AF555-EGF)-bound EGFR to Rab7-positive endosomes compared to control cells. However, TSG101 overexpression failed to restore the transport of ligand-bound EGFR to Rab7-positive endosomes in CPAP-depleted cells. Overall, these results, in the context of above-described observation that HRS overexpression restores TSG101 recruitment to endosomes in CPAP-depleted cells, suggest that Rab5-to-Rab7 conversion defects in CPAP-depleted cells are due to the failed TSG101 recruitment to EE.

**Fig. 9.**
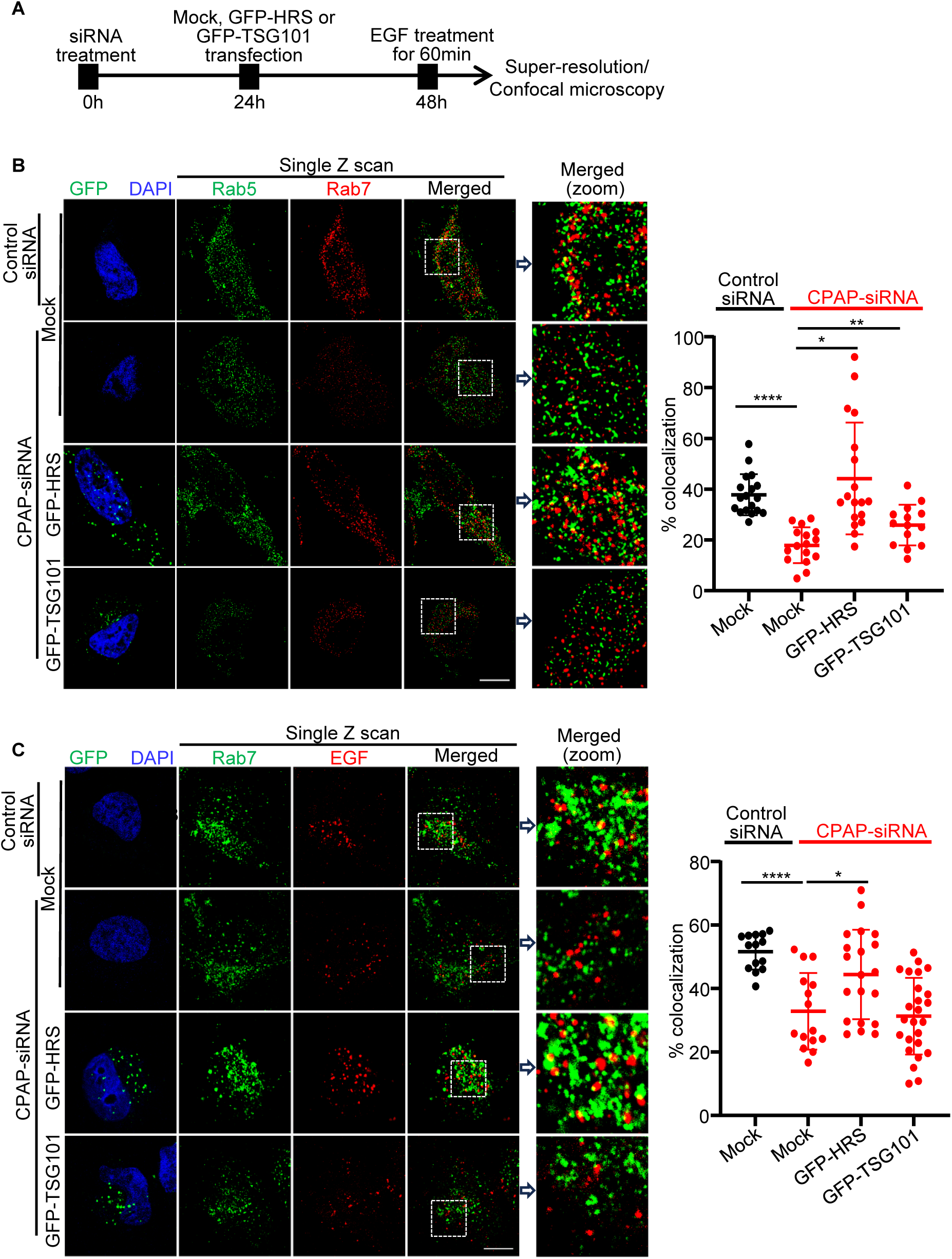
Rab5-to-Rab7 conversion and EGFR trafficking to late endosome are restored in CPAP-depleted cells upon HRS, but not TSG101, overexpression. A) Schematic depiction of experimental strategy using control- and CPAP-specific siRNA-treated HeLa cells with and without GFP-HRS or GFP-TSG101 expression. B&C) Control and CPAP-specific siRNA treated HeLa cells were subjected to mock or GFP-HRS or GFP-TSG101 vector transfection for 24h, treated with untagged EGF for 60 min and stained for Rab5 and Rab7 (B) or treated with AF555-EGF for 30 min and stained for Rab7 (C) and imaged by Lightning super resolution microscopy. Left panels: representative single Z-plane of images showing localization of Rab7 on Rab5-positive puncta (B) and AF555-EGF on Rab7-positive puncta (C) in cells with and without GFP-HRS or GFP-TSG101 expression. Right panels: colocalization (yellow) was quantified by counting percentages of Rab5-positive (green) puncta containing Rab7-positive (red) puncta in B and EGF-positive (red) puncta containing Rab7 (green, pseudo-color) in C in representative single Z-planes of each cell and quantified from multiple cells across at least 3 experiments. Zoomed images correspond to the dashed inset boxes of the indicated images. Scale bar: 10µm. *p*-values: *<0.05, ** <0.01, ****<0.0001 by unpaired non-parametric Mann-Whitney test.

### Rab5-to-Rab7 conversion and EGFR trafficking to MVB are restored in CPAP- and HRS-depleted cells upon exogenous CPAP expression

Finally, we examined whether restoring CPAP expression alone can restore Rab5-Rab7 conversion and EGFR-trafficking to MVBs in both CPAP- and HRS- depleted cells as shown in **Fig. 10A**. **Fig. 10B** shows that exogenous expression of siRNA-resistant CPAP resulted in a significant increase in localization of Rab7 on to the Rab5-positive endosomes during EVT progression in both CPAP- and HRS- depleted cells, compared to control cells, suggesting a restoration of Rab-to-Rab7 conversion. This data confirms that CPAP functions in parallel to HRS and mediates Rab5-to-Rab7 conversion during EVT. **Fig. 10C** shows that transport of ligand-bound EGFR to CD63-positive MVB also proceeded inconsequentially when CPAP expression was restored, again in both CPAP-and HRS-depleted cells. Overall, these results suggest that not only the disruptive effect of CPAP knockdown on TSG101 recruitment to the EE, Rab5-to-Rab7 conversion and endosome maturation during EVT is specific to lowered CPAP levels, but also the previously reported [17, 19], HRS-mediated recruitment of TSG101 on to endosomes for MVB formation can be compensated by CPAP.

**Fig. 10.**
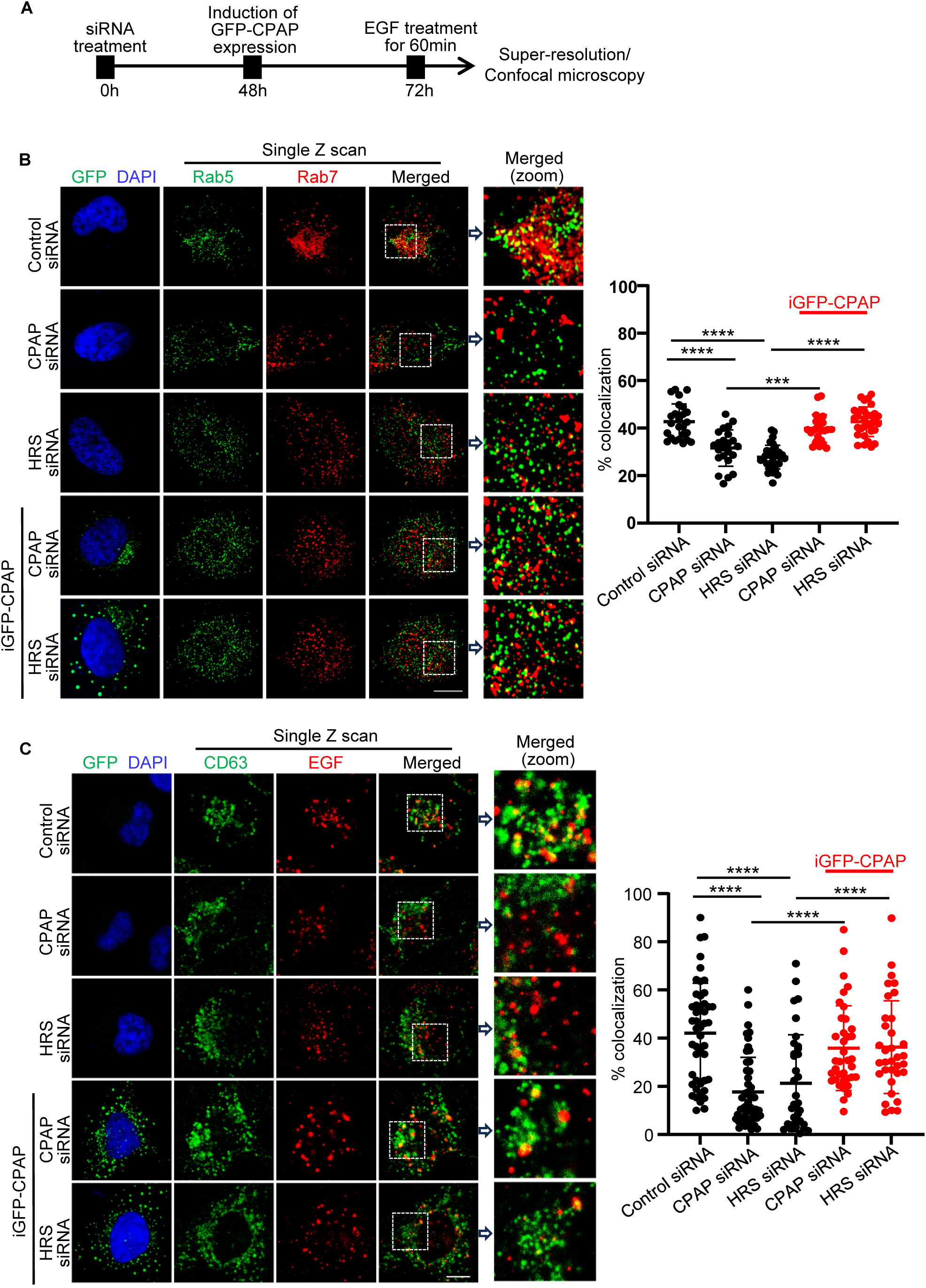
Rab5-to-Rab7 conversion and EGFR trafficking to MVB are restored in CPAP-and HRS-depleted cells upon exogenous CPAP expression. A) Schematic depiction of experimental strategy using control-, CPAP- and HRS-specific siRNA treated HeLa cells with and without siRNA-resistant GFP-CPAP expression. B) Airyscan super-resolution microscopy images showing cells stained for Rab5 and Rab7 at 60 min timepoint. Left panel: representative single Z-plane of images showing localization of Rab5- and Rab7-positive puncta in cells with and without GFP expression. Right panel: colocalization (yellow) was quantified by counting percentages of Rab5-positive (green) puncta containing Rab7-positive (red) puncta in representative single Z-planes of each cell and quantified from multiple cells across at least 3 experiments. C) Confocal microscopy images showing cells stained for CD63 and AF555-EGF at 60 min timepoint. Left panel: representative single Z-plane of images showing localization of CD63- and EGF-positive puncta in cells with and without GFP expression. Right panel: colocalization (yellow) was quantified by counting percentages of EGF-positive (red) puncta containing CD63-positive (green) puncta in representative single Z-planes of each cell and quantified from multiple cells across at least 3 experiments. Object-based colocalization macro tool of FIJI was employed for panels B and C. Zoomed images correspond to the dashed inset boxes of the indicated images. Scale bar: 10µm. *p*-values: ***<0.001, ****<0.0001 by unpaired non-parametric Mann-Whitney test.

Our data suggests that CPAP is an essential component of ESCRT, especially since endogenous HRS is unable to bypass CPAP function towards TSG101 recruitment during EVT. Our observations that CPAP and HRS overexpression can compensate for each other to recruit TSG101 on to early endosome suggest that CPAP and HRS function in parallel during MVB biogenesis and endosome maturation. This study reveals a previously unknown molecular role of the microcephaly- and centriole biogenesis-associated protein CPAP as an ESCRT pathway regulator.

## Discussion

The EVT, also referred to as endosomal trafficking, process is critical for many cellular functions, including regulation of signaling through cell surface receptors by targeting ligand bound receptors to the lysosome for degradation or recycling back to the plasma membrane. MVBs serve as critical intermediates during this process by sorting the endocytosed cargo and routing these cargo proteins to the lysosome [37]. In our recent reports [25], we have shown a novel positive regulatory role for CPAP, an essential centriole elongation protein [21, 24, 38], in MVB biogenesis and lysosome-targeting of ligand-bound EGFR and down-regulating signaling by this receptor. Here, we identify the molecular mechanism by which CPAP positively regulates MVB and lysosomal targeting of endocytosed cargo, wherein CPAP facilitates Rab5 to Rab7 conversion and ESCRT-I protein recruitment, both of which are critical steps for EE maturation to late endosomes/lysosomes.

Endosomal trafficking of the cell surface receptor cargo is tightly regulated through Rab GTPases such as Rab5 and Rab7. Progression from the EE to late endosome, which comprises of the intermediate stage MVB, also referred to as endosome maturation, has been shown to be regulated by a Rab5-to-Rab7 switch that involves an exchange between the cytosolic pool and endosomal membrane-bound fraction of Rab5 and Rab7 [15]. Completion of endosome maturation generates Rab7- and CD63-positive late endosomes/MVBs [39, 40]. Our previous report [25] showing defective EVT of endocytosed EGFR to late endosome and lysosome in CPAP-depleted cells suggested that Rab conversion and/or cellular levels and endosomal recruitment of Rab5 and Rab7 could be impacted upon CPAP depletion. In fact, we found that while localization of Rab5 to EE is comparable in control and CPAP-depleted cells, not only the Rab7 recruitment to endosomes is diminished, but also ligand-bound EGFR cargo failed to reach the Rab7-positive endosomes/lysosomes.

Importantly, Rab5-to-Rab7 conversion, which is indicated by the colocalization of these GTPases on endosomes, upon EVT induction using EGF, is severely diminished in CPAP-depleted cells. This observation suggested to us that CPAP may be regulating the cellular levels of Rab7 or being involved in recruiting this GTPase to endosome. However, our IB and IP studies showed that CPAP does not interact with either Rab5 or Rab7 and the cellular levels of these GTPases are also not considerably altered upon CPAP depletion suggesting that the suppressive effects of CPAP depletion on Rab5-to-Rab7 conversion, endosome maturation and receptor cargo transport are indirect effects on an upstream event.

Previous studies that used yeast and mammalian cells have shown that the evolutionarily conserved ESCRT-0, -I, -II, and -III complexes act sequentially in receptor sorting and MVB biogenesis. These ESCRT complexes are transiently recruited from the cytoplasm to endosome where they bind to internalized receptors that are marked for degradation by ubiquitination [41, 42]. HRS (ESCRT-0 protein) and TSG101 (ESCRT-I protein) are known to be functionally important in initiating the endosomal trafficking and MVB biogenesis process [17–19, 43]. HRS binds to TSG101 and mediates the targeting of ESCRT-I complex to EE [17]. On the other hand, TSG101 is critical for efficient degradation of EGFR, indicating its essential role in the sorting of cargo in MVBs and trafficking to lysosome [18] . ALIX, an ESCRT-associated accessory protein, is known to recruit ESCRT-III proteins to endosome during various ESCRT-dependent cellular processes including the ubiquitin-independent ILV sorting of G protein–coupled receptors PAR1 and P2Y1 for degradation [44, 45]. In this regard, although Rab5-to-Rab7 switch is also a critical step of endosome maturation, it is largely unknown how Rab conversion and ESCRT machinery functions are coordinated to generate MVB/late endosomes for trafficking of receptor cargo to lysosome. Since CPAP depletion diminished Rab conversion and endosome maturation, we studied if CPAP deficiency impacts EVT-induced endosome localization of key ESCRT proteins. Our data indicates that CPAP acts as a critical intermediary during the recruitment of TSG101 to endosome and potentially promotes Rab5-to-Rab7 conversion through ESCRT pathway function.

Although it has been reported that early- and late-acting ESCRTs preferentially localize on early and late endosomes respectively [46], it is now established that all ESCRTs can localize to EEA1- positive compartment during stimulation of endocytic vesicular transport [47, 48]. It has also been reported that HRS overexpression can cause accumulation of TSG101 in EE supporting the notion that HRS acts early and recruits TSG101 to EE membrane during EVT progression [18, 19]. Our results show that EGF stimulation of cells to induce EVT causes recruitment of HRS and TSG101, as well as the ESCRT-III associated protein ALIX to EE. Interestingly, our study has revealed, for the first time that similar to the above ESCRT proteins, CPAP is localized on to the EE in quiescent cells, and it is actively recruited to and colocalizes with ESCRT proteins on the endosome during EVT progression.

With respect to ESCRT recruitment, we observed that only TSG101, but not HRS or ALIX, localization on EE is disrupted in CPAP-depleted cells. Importantly, reintroduction of CPAP function restored TSG101 recruitment to the endosome in both CPAP and HRS-depleted cells suggesting that CPAP can compensate for HRS function, in terms of TSG101 recruitment to EE, during the vesicular transport. Furthermore, overexpression of the upstream ESCRT HRS, but not TSG101 itself, caused restoration of TSG101 localization on EE in CPAP-depleted cells. Overall, these observations suggest that CPAP acts in parallel to HRS for TSG101 recruitment to endosome during EVT and highlight the essential nature of CPAP function in EVT as endogenous HRS in CPAP-depleted cells fail to compensate for CPAP function, despite its intact cellular and endosomal levels.

Previous reports have shown that both HRS and TSG101 play key roles in MVB formation and cargo sorting for lysosomal targeting, with HRS being critical for recruiting downstream ESCRT components including TSG101 to endosome. On the other hand, TSG101 primarily functions by recognizing and binding ubiquitinated cargo proteins targeted for lysosomal degradation and promotes the stability of the vacuolar domains within the EE from which MVBs originate [18, 49]. Previous reports showing that depletion of HRS has only a modest effect on EGF degradation, compared with the almost complete inhibition of EGF degradation upon depletion of Tsg101 [18] not only validate this notion, but also suggest that MVB/late endosome formation are dependent on highly orchestrated functions of ESCRT components. Hence, our observations that TSG101 recruitment is disrupted in CPAP-depleted cells and restored by CPAP expression in HRS-depleted cells indicate that CPAP, at least in part, contributes to TSG101 recruitment to endosome and acts as an integral component of ESCRT machinery. Our results also show that the recruitment of ESCRT-III-associated protein ALIX to EE is independent of TSG101 recruitment.

HRS-TSG101 interaction has been reported before [17]. Our observations on the disrupted TSG101 localization to EE in CPAP-depleted cells suggested that CPAP interacts with TSG101 and/or other upstream ESCRT components and/or regulates the cellular levels of these proteins. In this regard, our results show that depletion of CPAP has no impact on the total cellular levels of HRS, TSG101 or ALIX suggesting that the disrupted TSG101 recruitment to endosome is not due to diminished protein levels. Lack of restoration of TSG101 recruitment to EE in CPAP-depleted cells, even upon overexpressing this protein substantiates this notion.

Importantly, our IP studies showing that CPAP can be immunoprecipitated using HRS-, TSG101- and ALIX-specific antibodies indicate that endogenous CPAP exists in physiological complexes which contain one or more ESCRT proteins. On the other hand, when exogenously expressed GFP-CPAP was immunoprecipitated, only TSG101, but not HRS or ALIX, was detected in the IP suggesting a potential direct interaction between CPAP and TSG101. Interestingly, EGF treatment to stimulate EVT of EGFR caused a profound increase in the amount of TSG101 bound to CPAP compared to unstimulated cells, albeit the amounts of GFP-CPAP in the IP were not considerably different, suggesting a possible interaction between CPAP and oligomerized TSG101. In fact, a previous report has shown that coiled-coil domain of TSG101 contributes to its homo-tetramerization [50].

Our observations that post-EGF-treatment, TSG101 pulldown by HRS and HRS pulldown by TSG101, are diminished under CPAP deficiency suggests that CPAP may be critical for a stable interaction between HRS and TSG101 during vesicular transport of receptor cargo. Previous reports [18, 51] have shown that TSG101 depletion inhibits MVB formation and alters the morphology of the EE in unstimulated cells, and more profoundly in EGF-stimulated cells suggesting that TSG101 is required for the formation of stable vacuolar domains within the EE that develop into MVBs. In this regard, our previous report on the diminished ILV budding/MVB formation and EGFR degradation under CPAP deficiency [25] and the current observations on CPAP-TSG101-HRS interaction dynamics propose the hypothesis that CPAP-TSG101 interaction is essential for the ESCRT function pertinent to MVB biogenesis during the vesicular transport of receptor cargo.

An important question of how the functional interaction between CPAP and ESCRT protein influences the Rab5-to-Rab7 conversion remains unanswered by our studies. A study employing yeast model has shown that ESCRT disruption inhibits the completion of Rab5-to-Rab7 conversion, but disruption of Rabs impedes ESCRT function, at endosomes as well [34] . Another study showed that ESCRTs function upstream of Rab conversion and loss of early ESCRTs leads to inhibition of Rab conversion [35]. Nevertheless, co-ordination between ESCRT and Rab conversion and the associated molecular events during vesicular transport are largely unknown. Our previous report [25] and current study together show that CPAP localizes to endosomes and depletion of CPAP function disrupts TSG101 (ESCRT-I) function, MVB formation and Rab conversion during EVT of EGFR. Consistent with TSG101 recruitment to endosomes, overexpression of ESCRT-0 protein HRS, but not TSG101 itself, caused effective restoration of Rab5-Rab7 conversion and EVT of ligand-bound EGFR to MVBs in CPAP-depleted cells. Our observations are consistent with the above-mentioned reports [34, 35] that suggest ESCRT functions upstream of Rab5-to-Rab7 conversion. Interestingly, CPAP overexpression in HRS-deficient cells also restored TSG101 recruitment to EE, Rab5-to-Rab7 conversion and trafficking of EGFR to late endosomes. Similarly, overexpression of, but not endogenous, HRS could compensate for CPAP in terms of recruitment of TSG101 to EE and endosome maturation. This clearly demonstrates that CPAP acts parallel to HRS during EVT and the CPAP-TSG101 axis is critical for Rab conversion.

In conclusion, we establish a previously unknown, non-centrosomal, endosome-associated function for a centriole biogenesis protein, CPAP. We show that CPAP facilitates ESCRT function and MVB biogenesis, Rab5-to-Rab7 conversion, and endosome maturation by recruiting an ESCRT-I member TSG101 to the EE during MVB formation. We also demonstrate, for the first time, CPAP localizes to EE and a functional interaction with ESCRT proteins on endosomes which is critical for bridging ESCRT-0 and ESCRT-I functions during the early events of MVB formation. Several centrosome-associated canonical cellular processes including centriole duplication, spindle orientation and positioning, and ciliogenesis have been attributed to CPAP before [22, 24, 26, 28, 29, 52]. Our recent reports have shown that CPAP has a role in positively regulating EVT of ligand-bound EGFR to lysosome for degradation to prevent persistent signaling by this receptor [25]. However, the role of CPAP in regulating ESCRT function and endosome maturation has never been described before.

Importantly, whether the newly identified, non-canonical function of CPAP in regulating ESCRT function and endosome maturation is required for the above-mentioned well-recognized canonical functions, or if they operate independently remains to be studied. Further studies are also required to establish if CPAP functions as a “bonafide” ESCRT component or a critical ESCRT-associated protein and how it is involved in various other ESCRT-dependent cellular functions. Nevertheless, our novel observations described here will open a new area of studies to determine if and how the fundamental cellular processes such as centriole duplication and endosomal transport of cargo proteins are coupled. Our observations on a critical role for CPAP in facilitating ESCRT function could also help to better explain the potential molecular mechanisms of CPAP-mutation associated diseases such as microcephaly.

## Resource availability

### 1. Lead contact

Resources for further information and resources should be directed to and will be fulfilled by the corresponding author, Dr. Radhika Gudi

### 2. Materials availability

This study did not generate new unique reagents.

### 3. Data availability

Any additional information required to reanalyze the data reported in this paper is available from the lead contact upon reasonable request.

## Acknowledgments

This work was supported by National Institutes of Health (NIH) grant R01DE030331 and R21DE026965 to C.V. and R.G. Unrestricted research funds from Hollings Cancer Center and College of Medicine, MUSC supported some aspects of the study. The authors are thankful for the Electron microscopy core and the Cell & Molecular Imaging Shared Resource which is supported by the Hollings Cancer Center, Medical University of South Carolina (P30 CA138313) and the Shared Instrumentation Grant S10 OD018113 as well as the flow cytometry core of MUSC for the FACS instrumentation support.

## Author contributions

R. G. conceived the idea, designed experiments, researched and analyzed data, and wrote/edited the manuscript, and C.V. designed experiments and wrote/edited the manuscript.

## Supplemental information

Document S1.

Figures S1-S13

## Materials and Methods

### Cell lines

HEK293T (National Gene Vector Biorepository) and HeLa (ATCC) cells were used in this study and cultured at 37°C. These cells were cultured in DMEM media, supplemented with 10% FBS, HEPES, sodium pyruvate, sodium bicarbonate and minimum essential amino acids. Cells were seeded for different transfection methods, based on manufacturer’s instructions or at approximately 50% confluency.

### Reagents

Transfection of HEK293T and HeLa cells using plasmids was done using calcium-phosphate method and Trans-X2 reagent (Mirus) respectively. Transfection of siRNA in both cell types was done by using the Trans-siQuest (Mirus). siRNA/shRNA regions targeted in CPAP have been described previously[25]. RNAi resistant GFP-CPAP overexpression construct was kindly provided by Dr. Pierre Gonczy (EPFL, Lausanne). The vector constructs expressing GFP-HRS and GFP-TSG101 were kindly provided by Dr. Clarisse Berlioz-Torrent (Institut Cochin, Paris). The vector constructs expressing GFP-Rab5, GFP-Rab11, and mCherry-Rab7, GFP-Rab7-WT and GFP-Rab7-DN were purchased from Addgene. siRNA against HRS, ALIX and TSG101 were purchased from Santa Cruz Biotech. Primary antibodies used in this study: rabbit anti-CPAP (Proteintech,11517-1-AP); mouse anti-CPAP (Abnova, H00055835-M01); anti-Rab5 (Proteintech, 20228-1-AP); anti-Rab7 (Proteintech, 55469-1-AP or Santa Cruz, sc-376362); Actin-HRP (Proteintech, 60008-1-Ig); -GFP (Proteintech, 66002-1-Ig or 50430-2-AP); -HRS (Santa Cruz, sc-271455); -TSG101 (Santa Cruz, sc-7964 ; Proteintech, 28283-1-AP); and -ALIX (Biolegend, 634502), EEA1 (Invitrogen, PA1-0337 or BD biosciences 610456) and CD63 (BD biosciences, 556019). Alexa fluor 555 (AF555)- tagged EGF and untagged EGF were purchased from Invitrogen and Tonbo biosciences respectively.

### Immunofluorescence

Cells were grown on sterile coverslips in a 24-well plate. After appropriate experimental treatments, cells were fixed with 4% paraformaldehyde. Permeabilization was done using 0.1% saponin-containing buffer for 30min at RT. Post-blocking with 1% BSA, primary and secondary antibody incubations were done at 37°C for 2h and 1h respectively. Coverslips were mounted using Prolong gold antifade mounting reagent (Invitrogen) and imaged using Zeiss 880 confocal or Airyscan or Leica Stellaris or Lightning super-resolution unit, using the 63X oil immersion objective with n.a. 1.4. Optimal recommended settings were used to acquire Z stacks.

### Image quantification and statistical considerations

Prior to image analysis, Airyscan files were automatically processed using the Zen software (version 2.0) of Zeiss (https://www.zeiss.com/microscopy/us/products/microscope-software/zen.html). Leica Lightning images were also obtained as processed and deconvolution-based images that were ready to be analyzed. Individual puncta were first delineated, and the background was removed by intensity thresholding in FIJI (https://imagej.nih.gov/ij/). Adjustments were made to entire images uniformly within an experiment, before using for quantification of colocalization. Images are presented as maximum intensity projection, or a single Z plane as indicated. Z-stack images were split into single Z planes and, for example: green and yellow pixels were quantified to determine the percentage of colocalization. Object-based colocalization was measured by counting all events of a single Z scan of each cell either manually or automatically by employing a macro function of FIJI (https://github.com/dsrichardson/fiji_macros/blob/master/2D_object_colocalization) as stated under each figure legend. A single Z-plane of each cell that showed maximum number of puncta was considered for quantification and image presentation. For manual counting, total number of channel 1 puncta (for example green channel) and those showing channel 2 puncta (for example red channel) colocalization (yellow) were counted for the representative single Z-plane of each cell individually using the multi-point tool and counter of FIJI for determining percentage colocalization. The FIJI macro, which is an object-based colocalization tool, identifies the center of mass of each vesicle (with fixed pixels in size for each experiment) in two channels of interest for colocalization analysis of percentage of channel 1 puncta on channel 2 puncta and vice versa. Find Maxima function was used for identifying the center of mass of each vesicle. Uniform maxima values and pixel sizes were used for automatic batch analysis of all images within a single experiment. Total fluorescence intensities of immunofluorescent stained cells were determined by calculating the integrated density using FIJI.

### Western blotting

Cells harvested at different time-points were lysed using RIPA buffer, supplemented with 0.1M PMSF and protease inhibitor cocktail. Lysates were then spun at 14000 rpm for 30min at 4°C, supernatants were separated and boiled at 95°C for 10m after addition of SDS sample buffer. Post-SDS-PAGE and Western blotting, immunoblotted PVDF membranes were developed using SuperSignal West Dura substrate (Thermo Scientific) chemiluminescence substrate and ChemiDoc imager (Biorad). Anti-rabbit-IgG HRP and -mouse-IgG HRP were obtained from Amersham and Biorad respectively. All blots are representative of at least three independent experiments with similar trends in results. Densitometry analysis was performed using FIJI and presented where appropriate.

### Transmission electron microscopy

HeLa cells treated with EGF ligand for 60min were fixed using 1% glutaraldehye, dehydrated and embedded in LR white resin (Ted Pella, Inc). 70nm thin sections on gold grids were treated with blocking buffer (20mM Hepes, 1% fish gelatin and 0.4% Triton X-100) at RT for 1h, incubated with rabbit anti-CPAP (Proteintech) or with mouse anti-EEA1 (BD biosciences) antibodies overnight at 4°C, followed by incubation with anti-rabbit IgG conjugated with 15nm-gold particles or anti-mouse IgG linked to 6nm-gold particles (Ted Pella, Inc) for 1h at RT. After extensive washing, sections were further stained with 1% Uranyl acetate for contrast and analyzed using JEOL1210 transmission electron microscope.

### Endosome fractionation

Cell pellets were rinsed twice with cold PBS and treated with 300 µl homogenization buffer (3mM imidazole, pH 7.4, 250 mM sucrose, 0.5 mM EDTA and protease inhibitors. Cells were homogenized with 20 strokes of a tight fitting Dounce homogenizer and centrifuged for 10 min at 3000 rpm in a tabletop centrifuge. The resulting post nuclear supernatant was adjusted to 40.6% sucrose using 62% sucrose. Diluted supernatant was placed in mini centrifuge tubes and overlayed with 700 μl of 35% and 500 μl of 25% sucrose solutions, followed by ultracentrifugation at 108,000g for 3h at 4°C. The endosome-enriched fraction at the 25/35% interface was collected, diluted threefold with cold PBS and centrifuged at 108,000g for 30 mins. Pellets were treated with SDS sample buffer and subjected to immunoblotting.

### Flow cytometry

Cells were fixed with 4% PFA for 15mins at RT, permeabilized with 1X permeabilization buffer (PB) containing 0.1% saponin for 30 mins at RT, blocked with 1% donkey serum in PB, incubated with primary antibodies for 2h at 37°C and fluorochrome-labeled secondary antibodies for 1h at 37°C. Cells were acquired using FACS Verse instrument and data was analyzed using the Flowjo software version 10.0 (https://www.flowjo.com/solutions/flowjo).

### EGFR internalization assay

HeLa cells were grown in serum free conditions overnight and incubated with media containing cycloheximide [CHX] (5μg/ml) for 1h at 37°C. Cells were treated in CHX media with EGF ligand (10ng/ml) for 1h on ice. Cells were washed with chilled serum free media and transferred to 37°C to induce internalization of receptor. For tracking routing of EGF receptor into vesicles by confocal, Alexa Fluor 555 conjugated EGF (250ng/ml) (Invitrogen) was used. Untagged EGF (10ng/ml) (Tonbo biosciences) was used for non-imaging or super-resolution imaging experiments.

### Statistical consideration

Experiments were repeated several times and all figure panels presented represent at least three experiments that produced similar trends in outcomes. Cumulative values or values from a representative experiment were used for graphical presentation and as indicated in figure legends. Each data point of graphs represents one cell and a single slice from each cell is used for counting all relevant channel puncta. Details on replicated and statistical significance are also mentioned in figure legends. *P*-values were calculated using GraphPad Prism statistical analysis software version 10 (https://www.graphpad.com/scientific-software/prism/). For most comparisons, unpaired non-parametric Mann-Whitney test was used. Densitometric values of IB from multiple experiments were compared by employing paired *t*-test.

## References

1. Scott, C.C., Vacca, F., and Gruenberg, J. (2014). Endosome maturation, transport and functions. Semin Cell Dev Biol 31, 2–10.

2. Schmid, S.L., Sorkin, A., and Zerial, M. (2014). Endocytosis: Past, present, and future. Cold Spring Harb Perspect Biol 6, a022509.

3. Vieira, A.V., Lamaze, C., and Schmid, S.L. (1996). Control of EGF receptor signaling by clathrin-mediated endocytosis. Science 274, 2086–2089.

4. Mettlen, M., Chen, P.H., Srinivasan, S., Danuser, G., and Schmid, S.L. (2018). Regulation of Clathrin-Mediated Endocytosis. Annual review of biochemistry 87, 871–896.

5. Neto, H., Collins, L.L., and Gould, G.W. (2011). Vesicle trafficking and membrane remodelling in cytokinesis. Biochem J 437, 13–24.

6. Mellman, I., and Nelson, W.J. (2008). Coordinated protein sorting, targeting and distribution in polarized cells. Nat Rev Mol Cell Biol 9, 833–845.

7. Villasenor, R., Kalaidzidis, Y., and Zerial, M. (2016). Signal processing by the endosomal system. Curr Opin Cell Biol 39, 53–60.

8. Tomas, A., Futter, C.E., and Eden, E.R. (2014). EGF receptor trafficking: consequences for signaling and cancer. Trends Cell Biol 24, 26–34.

9. Futter, C.E., Pearse, A., Hewlett, L.J., and Hopkins, C.R. (1996). Multivesicular endosomes containing internalized EGF-EGF receptor complexes mature and then fuse directly with lysosomes. J Cell Biol 132, 1011–1023.

10. Piper, R.C., and Katzmann, D.J. (2007). Biogenesis and function of multivesicular bodies. Annu Rev Cell Dev Biol 23, 519–547.

11. Gruenberg, J., and Stenmark, H. (2004). The biogenesis of multivesicular endosomes. Nat Rev Mol Cell Biol 5, 317–323.

12. Zerial, M. (1993). Regulation of endocytosis by the small GTP-ase rab5. Cytotechnology 11 Suppl 1, S47–49.

13. Stenmark, H. (2009). Rab GTPases as coordinators of vesicle traffic. Nature reviews. Molecular cell biology 10, 513–525.

14. Barbieri, M.A., Roberts, R.L., Gumusboga, A., Highfield, H., Alvarez-Dominguez, C., Wells, A., and Stahl, P.D. (2000). Epidermal growth factor and membrane trafficking. EGF receptor activation of endocytosis requires Rab5a. The Journal of cell biology 151, 539–550.

15. Rink, J., Ghigo, E., Kalaidzidis, Y., and Zerial, M. (2005). Rab conversion as a mechanism of progression from early to late endosomes. Cell 122, 735–749.

16. Poteryaev, D., Datta, S., Ackema, K., Zerial, M., and Spang, A. (2010). Identification of the switch in early-to-late endosome transition. Cell 141, 497–508.

17. Lu, Q., Hope, L.W., Brasch, M., Reinhard, C., and Cohen, S.N. (2003). TSG101 interaction with HRS mediates endosomal trafficking and receptor down-regulation. Proc Natl Acad Sci U S A 100, 7626–7631.

18. Razi, M., and Futter, C.E. (2006). Distinct roles for Tsg101 and Hrs in multivesicular body formation and inward vesiculation. Mol Biol Cell 17, 3469–3483.

19. Bache, K.G., Brech, A., Mehlum, A., and Stenmark, H. (2003). Hrs regulates multivesicular body formation via ESCRT recruitment to endosomes. J Cell Biol 162, 435–442.

20. Henne, W.M., Buchkovich, N.J., and Emr, S.D. (2011). The ESCRT pathway. Dev Cell 21, 77–91.

21. Cho, J.H., Chang, C.J., Chen, C.Y., and Tang, T.K. (2006). Depletion of CPAP by RNAi disrupts centrosome integrity and induces multipolar spindles. Biochem Biophys Res Commun 339, 742–747.

22. Hung, L.Y., Tang, C.J., and Tang, T.K. (2000). Protein 4.1 R-135 interacts with a novel centrosomal protein (CPAP) which is associated with the gamma-tubulin complex. Mol Cell Biol 20, 7813–7825.

23. Hung, L.Y., Chen, H.L., Chang, C.W., Li, B.R., and Tang, T.K. (2004). Identification of a novel microtubule-destabilizing motif in CPAP that binds to tubulin heterodimers and inhibits microtubule assembly. Mol Biol Cell 15, 2697–2706.

24. Tang, C.J., Fu, R.H., Wu, K.S., Hsu, W.B., and Tang, T.K. (2009). CPAP is a cell-cycle regulated protein that controls centriole length. Nat Cell Biol 11, 825–831.

25. Gudi, R., Palanisamy, V., and Vasu, C. (2021). Centrosomal P4.1-associated protein (CPAP) positively regulates endocytic vesicular transport and lysosome targeting of EGFR. Sci Rep 11, 12689.

26. Kohlmaier, G., Loncarek, J., Meng, X., McEwen, B.F., Mogensen, M.M., Spektor, A., Dynlacht, B.D., Khodjakov, A., and Gonczy, P. (2009). Overly long centrioles and defective cell division upon excess of the SAS-4-related protein CPAP. Curr Biol 19, 1012–1018.

27. Schmidt, T.I., Kleylein-Sohn, J., Westendorf, J., Le Clech, M., Lavoie, S.B., Stierhof, Y.D., and Nigg, E.A. (2009). Control of centriole length by CPAP and CP110. Curr Biol 19, 1005–1011.

28. Wu, K.S., and Tang, T.K. (2012). CPAP is required for cilia formation in neuronal cells. Biol Open 1, 559–565.

29. Zheng, X., Ramani, A., Soni, K., Gottardo, M., Zheng, S., Ming Gooi, L., Li, W., Feng, S., Mariappan, A., Wason, A., et al. (2016). Molecular basis for CPAP-tubulin interaction in controlling centriolar and ciliary length. Nat Commun 7, 11874.

30. Gabriel, E., Wason, A., Ramani, A., Gooi, L.M., Keller, P., Pozniakovsky, A., Poser, I., Noack, F., Telugu, N.S., Calegari, F., et al. (2016). CPAP promotes timely cilium disassembly to maintain neural progenitor pool. EMBO J 35, 803–819.

31. Sharma, A., Aher, A., Dynes, N.J., Frey, D., Katrukha, E.A., Jaussi, R., Grigoriev, I., Croisier, M., Kammerer, R.A., Akhmanova, A., et al. (2016). Centriolar CPAP/SAS-4 Imparts Slow Processive Microtubule Growth. Dev Cell 37, 362–376.

32. Gudi, R.R., Janakiraman, H., Howe, P.H., Palanisamy, V., and Vasu, C. (2021). Loss of CPAP causes sustained EGFR signaling and epithelial-mesenchymal transition in oral cancer. Oncotarget 12, 807–822.

33. Babst, M. (2011). MVB vesicle formation: ESCRT-dependent, ESCRT-independent and everything in between. Curr Opin Cell Biol 23, 452–457.

34. Russell, M.R., Shideler, T., Nickerson, D.P., West, M., and Odorizzi, G. (2012). Class E compartments form in response to ESCRT dysfunction in yeast due to hyperactivity of the Vps21 Rab GTPase. J Cell Sci 125, 5208–5220.

35. Ott, D.P., Desai, S., Solinger, J.A., Kaech, A., and Spang, A. (2025). Coordination between ESCRT function and Rab conversion during endosome maturation. EMBO J 44, 1574–1607.

36. Pornillos, O., Higginson, D.S., Stray, K.M., Fisher, R.D., Garrus, J.E., Payne, M., He, G.P., Wang, H.E., Morham, S.G., and Sundquist, W.I. (2003). HIV Gag mimics the Tsg101-recruiting activity of the human Hrs protein. J Cell Biol 162, 425–434.

37. Woodman, P.G., and Futter, C.E. (2008). Multivesicular bodies: co-ordinated progression to maturity. Curr Opin Cell Biol 20, 408–414.

38. Kleylein-Sohn, J., Westendorf, J., Le Clech, M., Habedanck, R., Stierhof, Y.D., and Nigg, E.A. (2007). Plk4-induced centriole biogenesis in human cells. Dev Cell 13, 190–202.

39. Vitelli, R., Santillo, M., Lattero, D., Chiariello, M., Bifulco, M., Bruni, C.B., and Bucci, C. (1997). Role of the small GTPase Rab7 in the late endocytic pathway. J Biol Chem 272, 4391–4397.

40. Ceresa, B.P., and Bahr, S.J. (2006). rab7 activity affects epidermal growth factor:epidermal growth factor receptor degradation by regulating endocytic trafficking from the late endosome. J Biol Chem 281, 1099–1106.

41. Shields, S.B., and Piper, R.C. (2011). How ubiquitin functions with ESCRTs. Traffic 12, 1306–1317.

42. Korbei, B. (2022). Ubiquitination of the ubiquitin-binding machinery: how early ESCRT components are controlled. Essays Biochem 66, 169–177.

43. Raiborg, C., Bache, K.G., Mehlum, A., Stang, E., and Stenmark, H. (2001). Hrs recruits clathrin to early endosomes. EMBO J 20, 5008–5021.

44. Dores, M.R., Paing, M.M., Lin, H., Montagne, W.A., Marchese, A., and Trejo, J. (2012). AP-3 regulates PAR1 ubiquitin-independent MVB/lysosomal sorting via an ALIX-mediated pathway. Mol Biol Cell 23, 3612–3623.

45. Bissig, C., and Gruenberg, J. (2014). ALIX and the multivesicular endosome: ALIX in Wonderland. Trends Cell Biol 24, 19–25.

46. Frankel, E.B., and Audhya, A. (2018). ESCRT-dependent cargo sorting at multivesicular endosomes. Semin Cell Dev Biol 74, 4–10.

47. Mayers, J.R., and Audhya, A. (2012). Vesicle formation within endosomes: An ESCRT marks the spot. Commun Integr Biol 5, 50–56.

48. Slagsvold, T., Pattni, K., Malerod, L., and Stenmark, H. (2006). Endosomal and non-endosomal functions of ESCRT proteins. Trends Cell Biol 16, 317–326.

49. Sundquist, W.I., Schubert, H.L., Kelly, B.N., Hill, G.C., Holton, J.M., and Hill, C.P. (2004). Ubiquitin recognition by the human TSG101 protein. Mol Cell 13, 783–789.

50. White, J.T., Toptygin, D., Cohen, R., Murphy, N., and Hilser, V.J. (2017). Structural Stability of the Coiled-Coil Domain of Tumor Susceptibility Gene (TSG)-101. Biochemistry 56, 4646–4655.

51. Doyotte, A., Russell, M.R., Hopkins, C.R., and Woodman, P.G. (2005). Depletion of TSG101 forms a mammalian “Class E” compartment: a multicisternal early endosome with multiple sorting defects. J Cell Sci 118, 3003–3017.

52. Kitagawa, D., Kohlmaier, G., Keller, D., Strnad, P., Balestra, F.R., Fluckiger, I., and Gonczy, P. (2011). Spindle positioning in human cells relies on proper centriole formation and on the microcephaly proteins CPAP and STIL. J Cell Sci 124, 3884–3893.

