## Supplemental Figures for "Centrosomal P4.1-associated protein (CPAP) is a novel regulator of ESCRT pathway function during endosome maturation"

Fig. S1

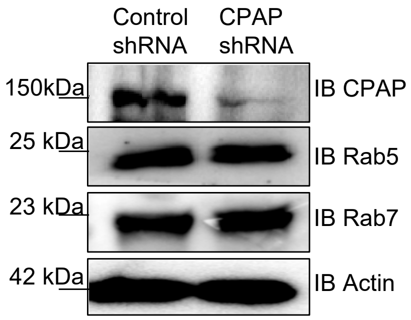


**Supplemental Fig. 1: Cellular levels of Rab5 and Rab7 are not impacted by CPAP depletion.** Immunoblotting of control and CPAP shRNA-expressing HeLa cell lysates was performed with rabbit anti-CPAP antibody, and anti-Rab5 and anti-Rab7 antibodies.

Fig. S2

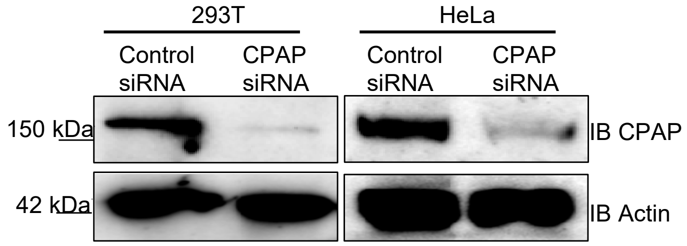


**Supplemental Fig. 2: Validating CPAP-specific antibody**. Lysates of HEK293T and HeLa cells treated with control or CPAP siRNA (for 48 hours) were immunoblotted with mouse anti-CPAP antibody.

Fig. S3

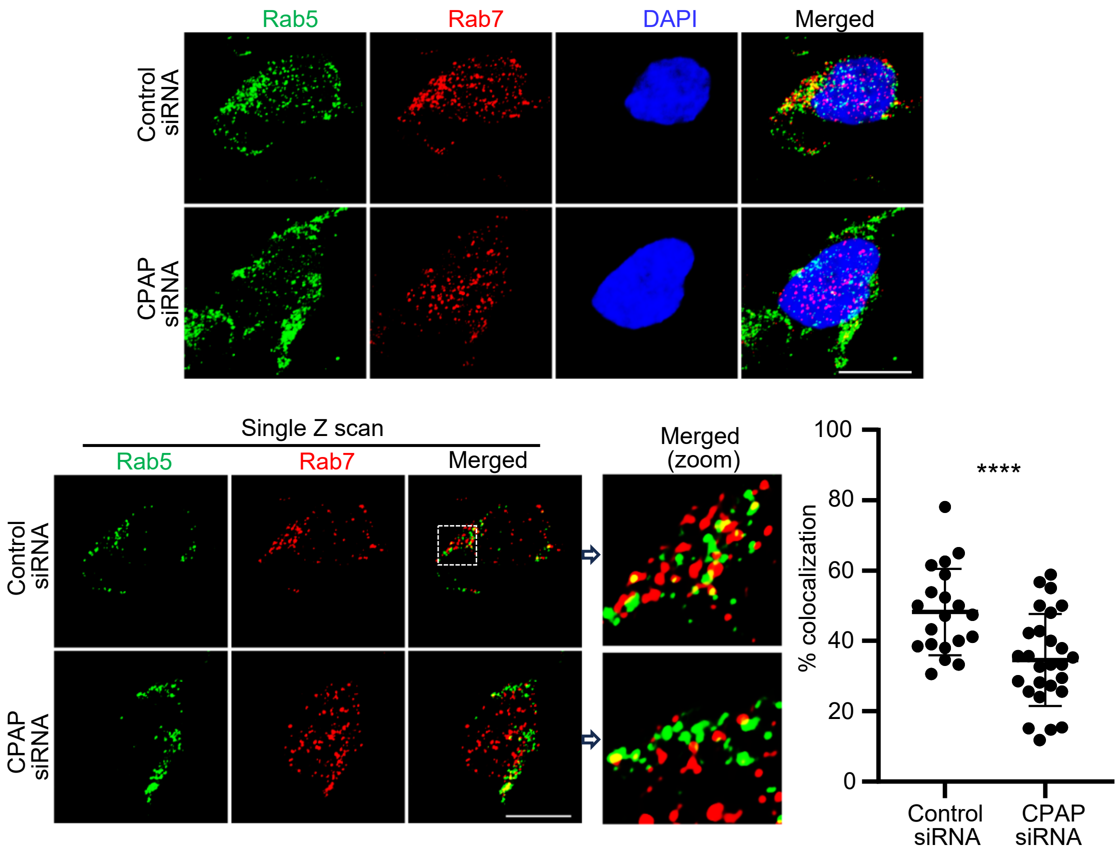


**Supplemental Fig. 3: Rab5-to-Rab7 conversion is hampered in CPAP siRNA-treated cells.** HeLa cells were treated with control- or CPAP-siRNA for 48 hours and incubated with untagged EGF for 60 minutes and imaged by Airyscan super-resolution microscopy. Top panel: maximum intensity-projection images of Z stacks; bottom left panel: single Z-plane of images; bottom right panel: colocalization (yellow) was quantified by counting percentages of Rab5-positive (green) puncta containing Rab7 (red) puncta in representative single Z-planes of each cell and quantified from multiple cells across at least 3 experiments. Object-based colocalization macro tool of FIJI was employed. Scale bar: 10µm. *p*-value: ****<0.0001 by unpaired non-parametric Mann-Whitney test.

Fig. S4

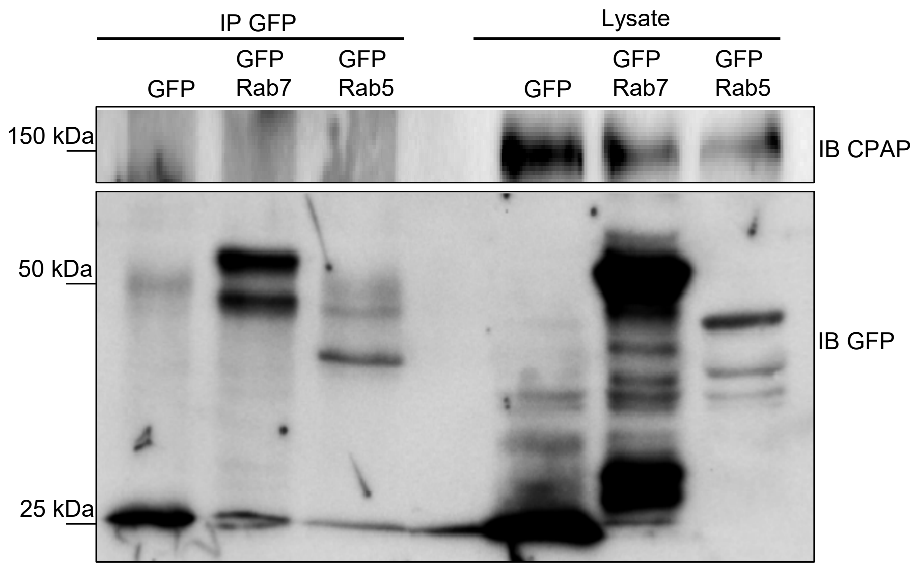


**Supplemental Fig. 4: CPAP was not detected in Rab5- and Rab7- specific IPs.** HEK293T cells transfected with GFP or GFP-Rab7 and GFP-Rab5 expression vectors were subjected to immunoprecipitation with the anti-GFP antibody. Immunoblotting was performed as indicated.

Fig. S5

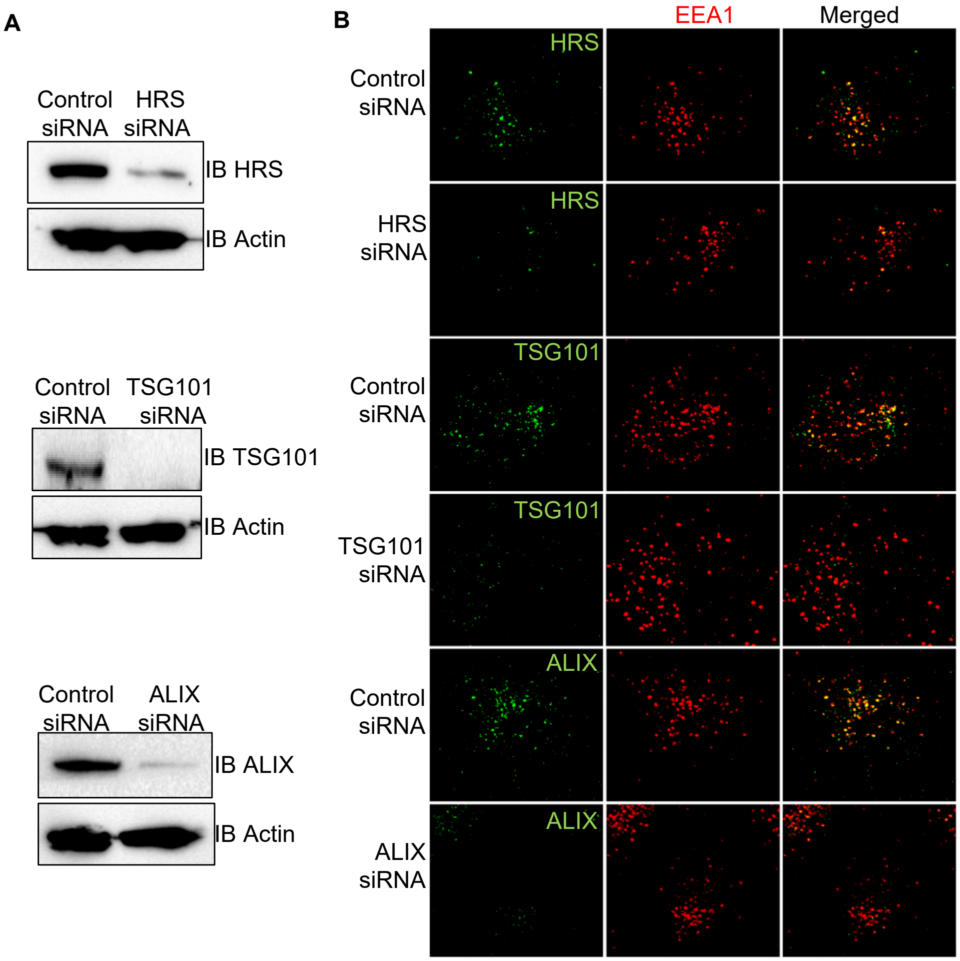


**Supplemental Fig. 5: Validating ESCRT protein-specific antibodies.** Control or HRS, TSG101, and ALIX specific siRNA treated HeLa cells (for 48 hours) were tested for knockdown efficiency by immunoblotting (left panels) as well as by Airyscan super-resolution microscopy (right panels). Right panel shows single Z images of representative cells.

Fig. S6

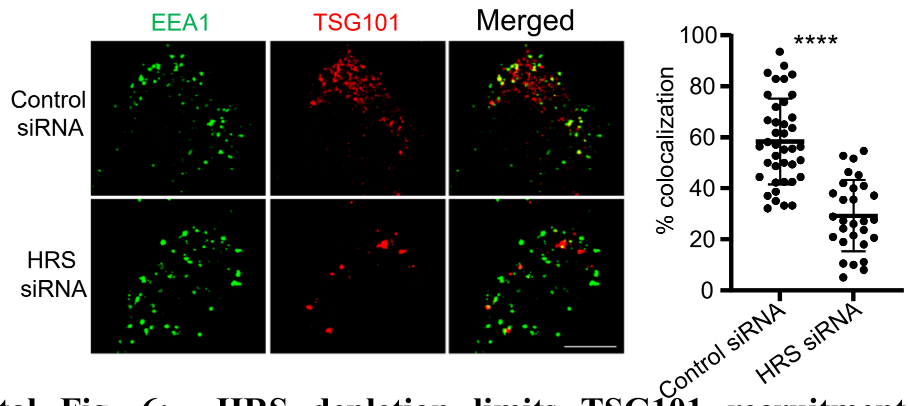


**Supplemental Fig. 6: HRS depletion limits TSG101 recruitment to the early endosome during EVT.** Control or HRS-siRNA treated HeLa cells were exposed to EGF for 60 minutes and stained using anti-EEA1 and -TSG101 antibodies. Images were acquired by Airyscan super-resolution microscopy. Left panel: representative single Z-plane of images showing localization of TSG101 on EEA1-positive puncta. Right panel: colocalization (yellow) was quantified by counting percentages of EEA1-positive (green) puncta containing TSG101 (red) puncta in representative single Z-planes of each cell and quantified from multiple cells across at least 3 experiments. Scale bar: 10µm. *p*-value: ****<0.0001 by unpaired non-parametric Mann-Whitney test.

Fig. S7

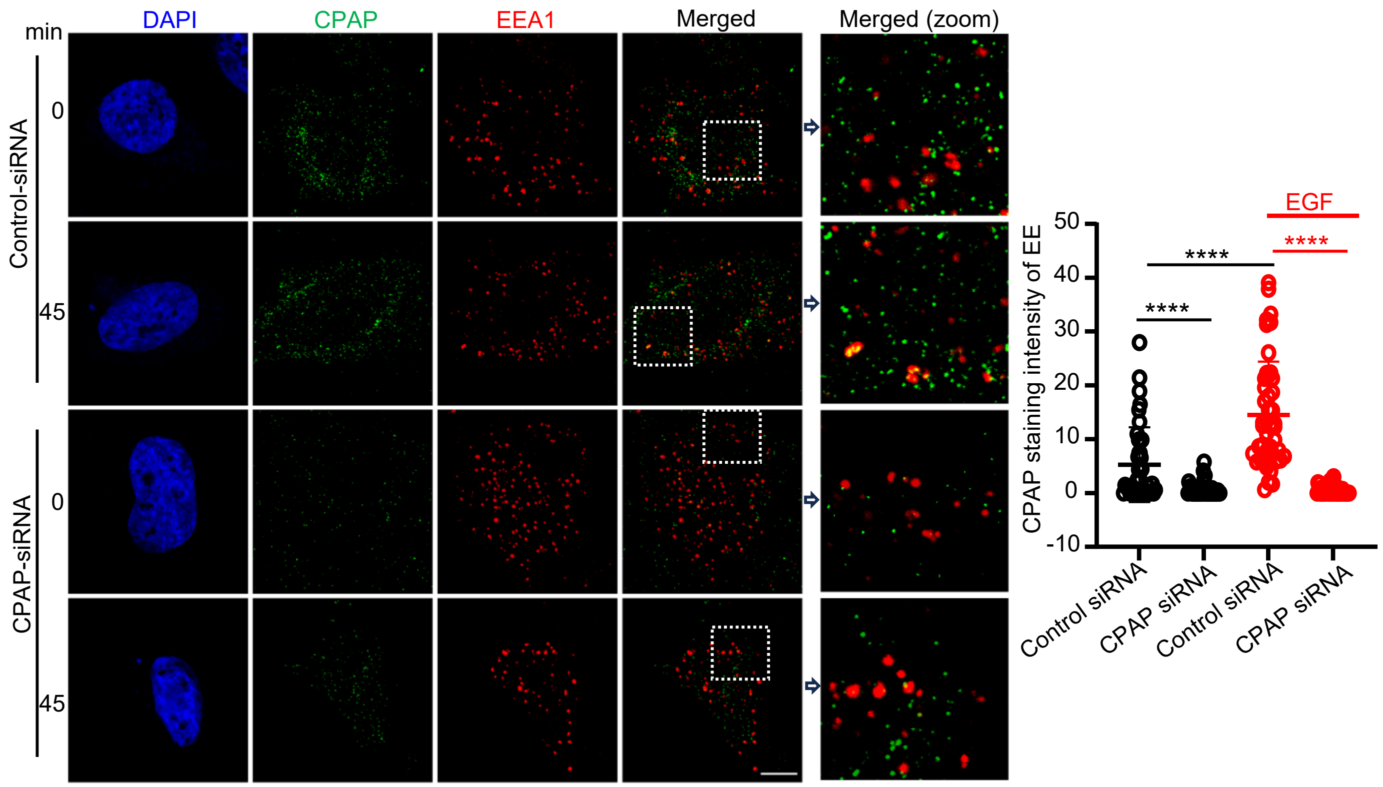


**Supplemental Fig. 7: CPAP is detected on EE in quiescent and EGF-treated cells.** Control-(top panel) or CPAP-siRNA-(bottom panel) treated HeLa cells were left untreated (0 minutes) or treated with EGF for 45 minutes, followed by staining with rabbit anti-CPAP and -mouse anti-EEA1 antibodies. Images were acquired using Lightning super-resolution microscope. Relative integrated fluorescence intensity values of CPAP staining on individual EEA1-positive endosome was quantified in representative single Z-planes of each cell and quantified from multiple cells. Zoomed images correspond to the dashed inset boxes of the indicated images. Scale bar: 10µm. *p*-value: ****<0.0001 by unpaired non-parametric Mann-Whitney test.

Fig. S8

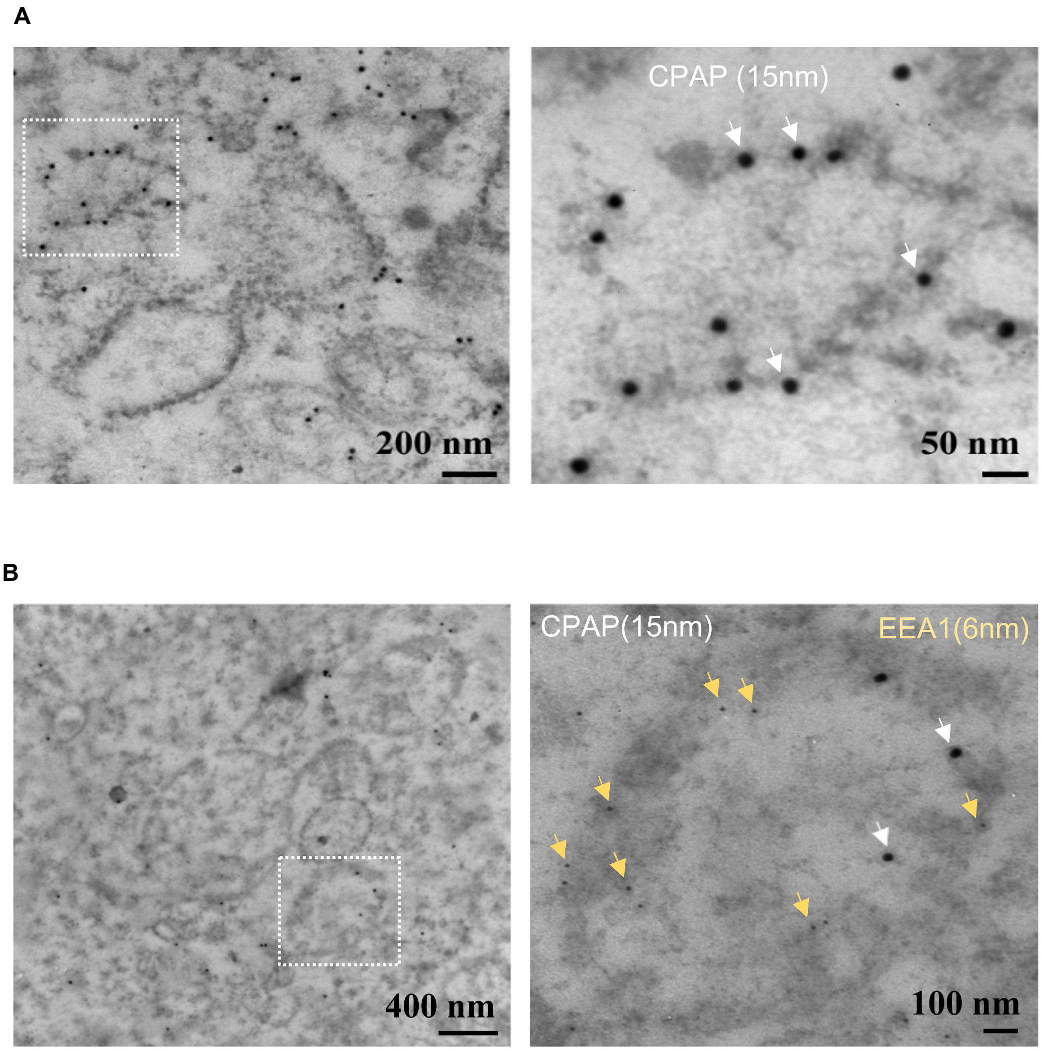


**Supplemental Fig. 8: CPAP expression is observed on endosomes by TEM.** HeLa cells treated with EGF for 60 minutes were subjected to transmission electron microscopy (TEM)-immunogold analysis. 70nm thin sections on gold grids were stained with rabbit anti-CPAP antibody alone (A) or along with -mouse anti-EEA1 antibody (B), and anti-rabbit-IgG-gold (15nm) and anti-mouse IgG-gold (6nm) conjugates.

Fig. S9

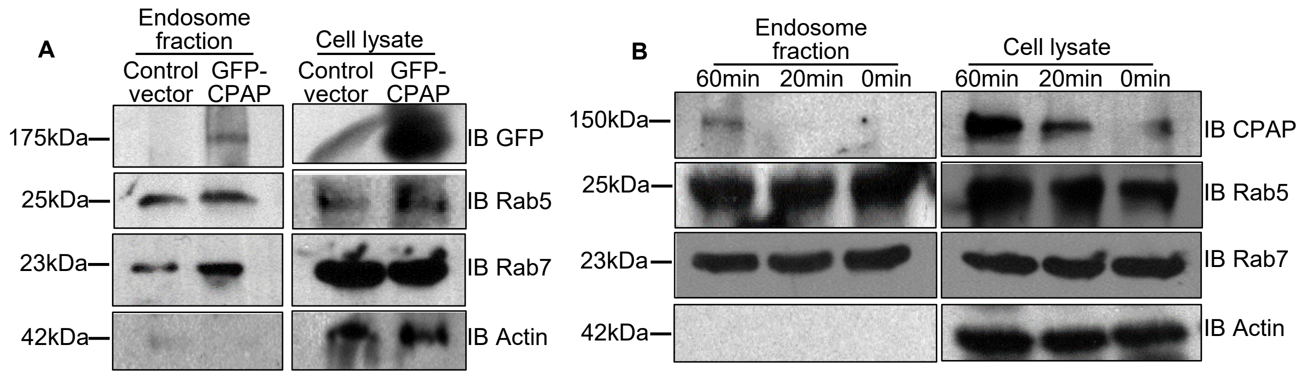


**Supplemental Fig. 9: CPAP expression is detected in endosome-fractions.** A) HEK293T cells stably expressing doxycycline inducible GFP-CPAP were subjected to endosome fractionation by ultracentrifugation approach. Endosome fraction and whole cell lysates as input control were probed for indicated proteins by IB. B) HEK293T cells left untreated or treated with untagged EGF for different time points were subjected to endosome fractionation, and the endosome fractions and whole cell lysate were subjected to IB for indicated protein.

Fig. S10

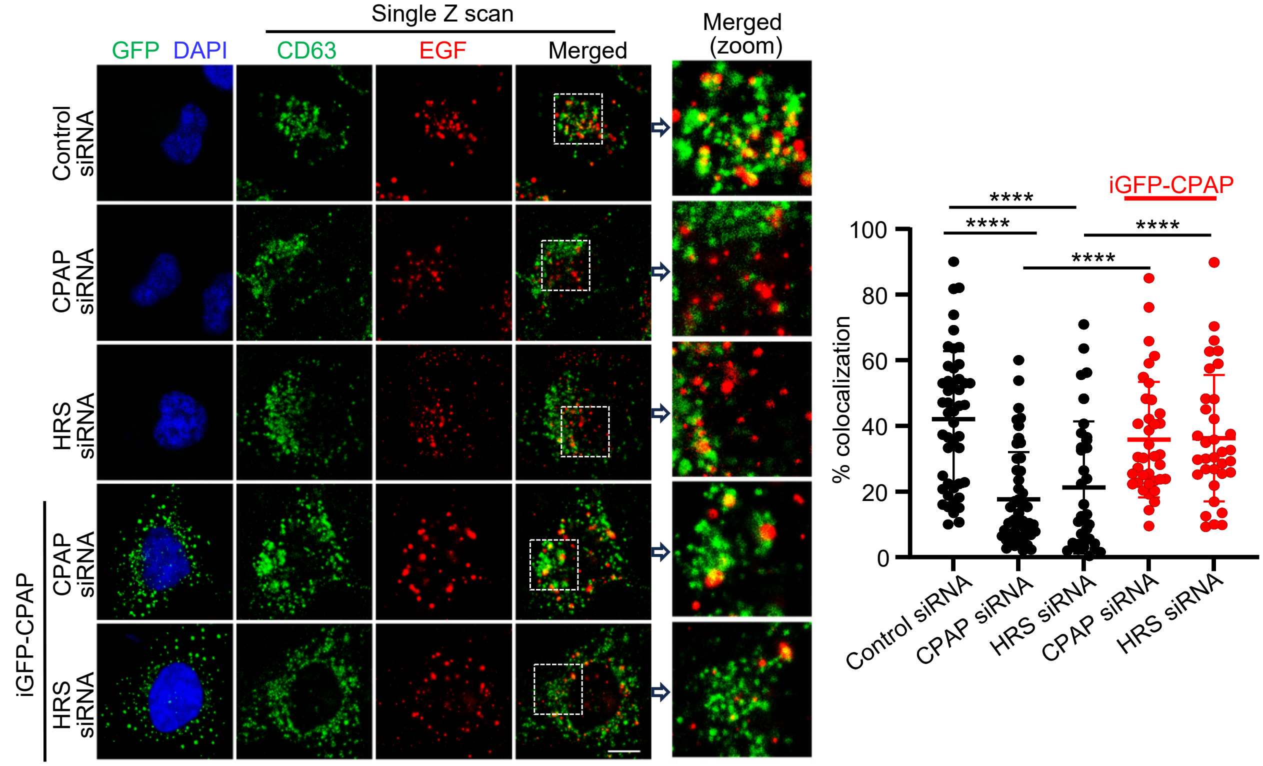


**Supplemental Fig.10: Exogenous expression of CPAP restores EVT function in CPAP- and HRS-depleted cells.** Cells were treated similar to Fig. 6A. AF555-linked EGF- treated cells were stained for CD63 and imaged using the confocal. Left panel: representative single Z-plane of images showing localization of AF555-EGF on CD63-positive puncta in cells treated as indicated. Right panel: colocalization (yellow) was quantified by counting percentages of EGF-positive (red) puncta containing CD63 (green) puncta in representative single Z-planes of each cell and quantified from multiple cells across at least 3 experiments. Scale bar: 10μm. *p*-value: ****<0.0001 by unpaired non-parametric Mann-Whitney test.

Fig. S11

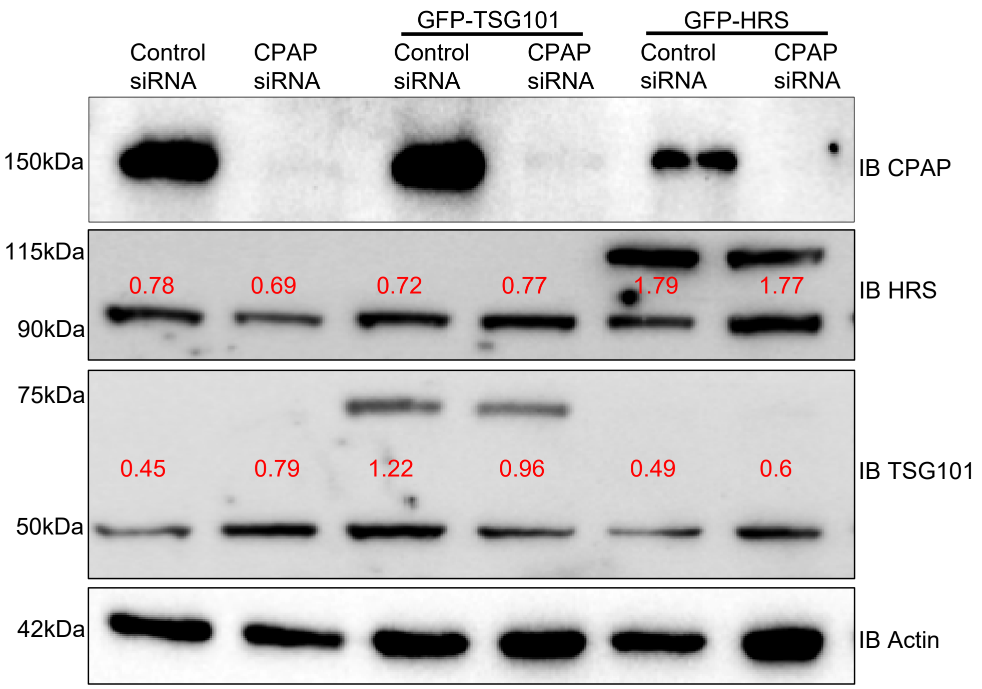


**Supplemental Fig. 11: Exogenous expression of GFP-TSG101 and GFP-HRS in control- and CPAP-siRNA cells.** Lysates of control- or CPAP-specific siRNA-treated HEK293T cells transfected with either GFP-TSG101 or GFP-HRS were subjected to immunoblotting with indicated antibodies. Densitomety values obtained for each lane using FIJI software are shown for relevant groups.

Fig. S12

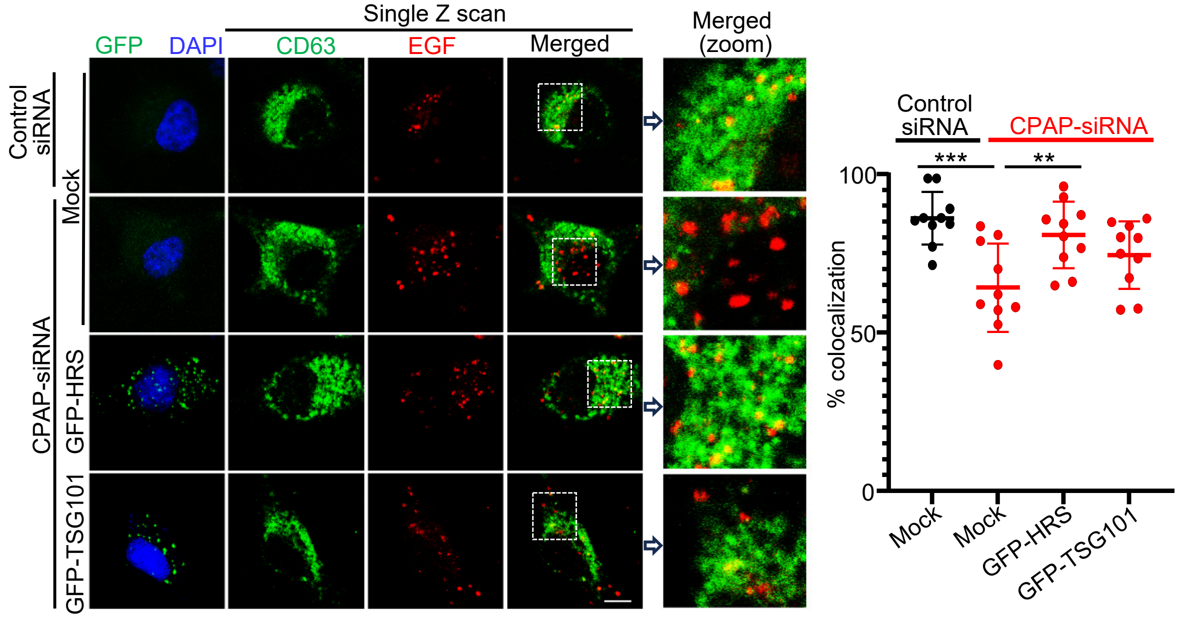


**Supplemental Fig. 12: HRS overexpression restored EGFR trafficking to MVB in CPAP-depleted cells.** Similar to Fig. 9C, cells treated with AF555-linked EGF were stained for CD63 and imaged as done earlier. Left panel: representative single Z-plane of images showing localization of AF555-EGF on CD63 positive puncta in cells with and without GFP-HRS or GFP-TSG101 expression. Right panel: percentages of CD63 (green, pseudo-color) puncta showing EGF (red) colocalization (yellow) quantified in single Z-planes of multiple cells across at least 3 experiments. Scale bar: 10µm. *p*-values: *<0.05, ** <0.01, ****<0.0001.

Fig. S13

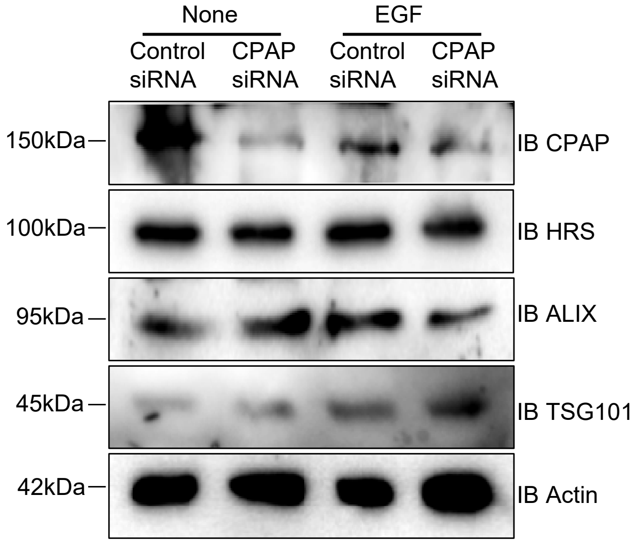


**Supplemental Fig. 13: Cellular levels of ESCRT proteins are unchanged upon EGF treatment.** Control- or CPAP-siRNA treated HEK293T cells were treated with EGF for 60 min and subjected to immunoblotting with indicated ESCRT-protein specific antibodies and anti-CPAP antibody.
